# Structural basis for substrate selection by the SARS-CoV-2 replicase

**DOI:** 10.1101/2022.05.20.492815

**Authors:** Brandon F. Malone, Jason K. Perry, Paul Dominic B. Olinares, James Chen, Todd K. Appelby, Joy Y. Feng, John P. Bilello, Honkit Ng, Johanna Sotiris, Mark Ebrahim, Eugene Y.D. Chua, Joshua H. Mendez, Ed T. Eng, Robert Landick, Brian T. Chait, Elizabeth A. Campbell, Seth A. Darst

## Abstract

The SARS-CoV-2 RNA-dependent RNA polymerase coordinates viral RNA synthesis as part of an assembly known as the replication-transcription complex (RTC)^1^. Accordingly, the RTC is a target for clinically approved antiviral nucleoside analogs, including remdesivir^2^. Faithful synthesis of viral RNAs by the RTC requires recognition of the correct nucleotide triphosphate (NTP) for incorporation into the nascent RNA. To be effective inhibitors, antiviral nucleoside analogs must compete with the natural NTPs for incorporation. How the SARS-CoV-2 RTC discriminates between the natural NTPs, and how antiviral nucleoside analogs compete, has not been discerned in detail. Here, we use cryo-electron microscopy to visualize the RTC bound to each of the natural NTPs in states poised for incorporation. Furthermore, we investigate the RTC with the active metabolite of remdesivir, remdesivir triphosphate (RDV-TP), highlighting the structural basis for the selective incorporation of RDV-TP over its natural counterpart ATP^3,4^. Our results elucidate the suite of interactions required for NTP recognition, informing the rational design of antivirals. Our analysis also yields insights into nucleotide recognition by the nsp12 NiRAN, an enigmatic catalytic domain essential for viral propagation^5^. The NiRAN selectively binds GTP, strengthening proposals for the role of this domain in the formation of the 5’ RNA cap^6^.

---

COVID-19, caused by the coronavirus SARS-CoV-2, continues to devastate livelihoods, and overwhelm healthcare systems worldwide. Given the urgency of stymieing infection and mitigating disease morbidity, a concerted research effort has begun to elucidate the molecular details of the viral lifecycle and to design therapeutics to disrupt it ^1^. The viral RNA-dependent RNA polymerase (RdRp, encoded by non-structural protein 12, or nsp12) functions as part of the holo-RdRp (comprising the RdRp and auxiliary proteins nsp7 and nsp8 as a heterotetramer nsp7/nsp8_2_/nsp12) in a replication–transcription complex (holo-RdRp + RNA, or RTC) to direct all viral RNA synthesis in conjunction with a coterie of viral nucleic acid processing enzymes. In less than two years, targeting the RTC by the antivirals remdesivir (RDV) and molnupiravir has become the staple of clinical care ^2,7^. This clinical success underscores the importance of the RTC as a pharmacological target and incentivizes the design of more efficacious nucleotide analogues for the treatment of COVID-19. Furthermore, due to the highly conserved nature of the coronavirus RdRp active site ^8^, nucleotide analogue inhibitors of the RTC are excellent candidates for pan-viral inhibitors that would be effective against emerging variants and may be repurposed to tackle future pathogenic coronaviruses that could arise.

Biochemical and single-molecule experiments have characterized the RNA synthesis activity of the SARS-CoV-2 RTC, its selectivity for a wide range of nucleotide analogs, and their effects on RNA synthesis ^3,4,9,10^. These investigations yielded insights into the ability of the RTC to discriminate between natural NTPs and related antiviral analogs. Studies of remdesivir (RDV) revealed that the active metabolite of RDV, RDV triphosphate (RDV-TP), possesses enhanced selectivity for incorporation into nascent RNA over its natural counterpart, ATP ^3,4^. Rapid, competitive incorporation is likely a critical facet of the clinical effectiveness of RDV. However, the structural basis for NTP recognition and RDV-TP selectivity remains unknown, necessitating structural examinations of stalled RTC ternary complexes with incoming NTP substrates and catalytic metal ions poised for catalysis (Michaelis or pre-incorporation complexes) (Fig. 1a).

**Fig. 1.**
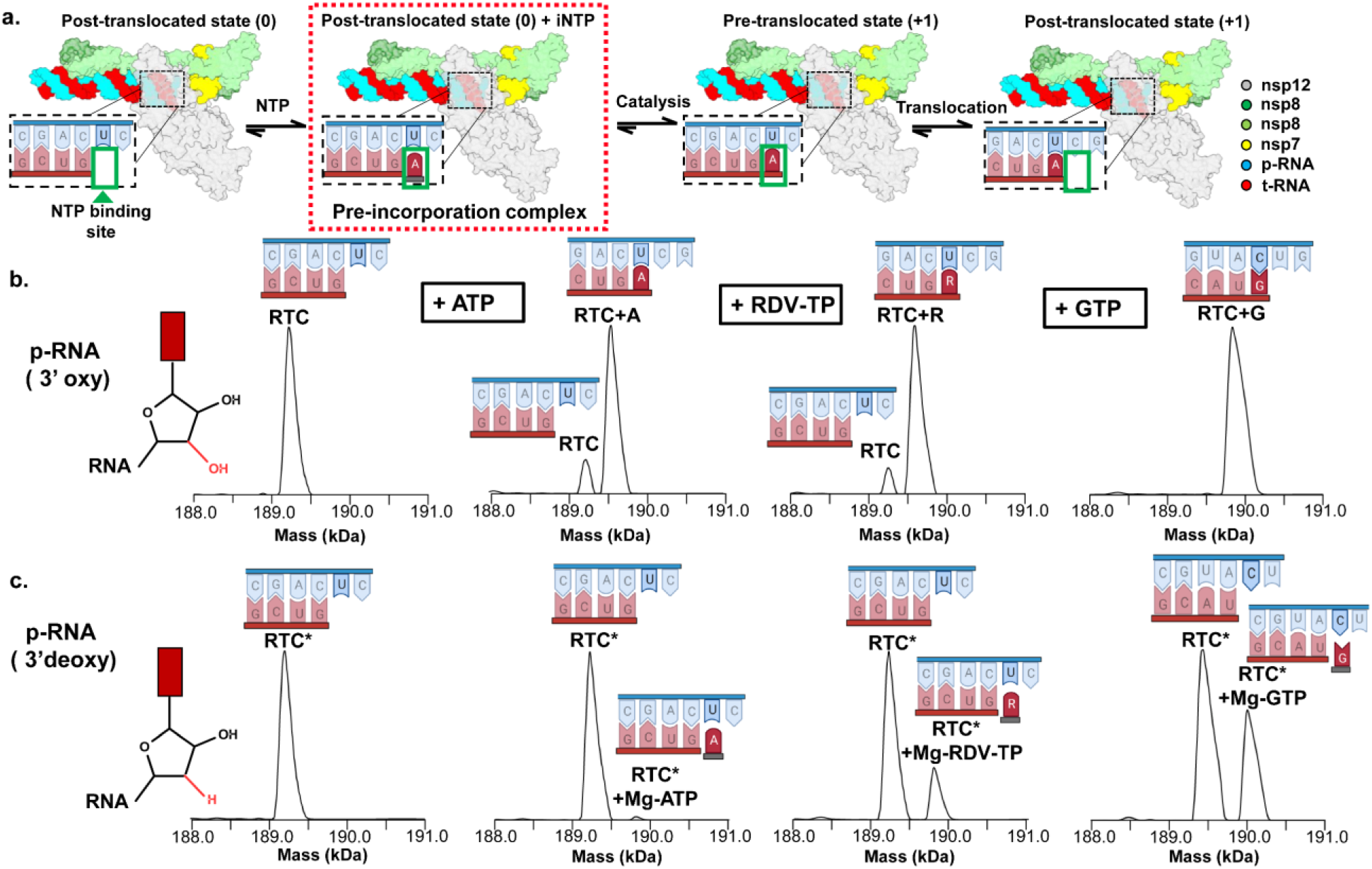
Capturing the SARS-CoV-2 RTC ternary complex. **(a)** Schematic depicting the major steps of the nucleotide addition cycle of the replication-transcription complex. The pre-incorporation complex studied here is highlighted (red dashed box). **(b)** Native mass spectrometry (nMS) analysis of the RTC bound to a 3’oxy product-RNA (p-RNA) in the absence and presence of 300 μM ATP, RDV-TP, or GTP respectively. **(c)** Similar nMS analysis to **(b)** but the RTC was reconstituted using a 3’-deoxy p-RNA (RTC*).

Stepwise nucleotide incorporation necessitates transferring the 3’-end of the product-RNA (p-RNA) to the post-translocated site, allowing binding of the incoming NTP substrate along with two catalytic metal ions in the active-site cleft (Fig. 1a) ^11,12,13^. In the subsequent catalytic step(s), the geometry of correct Watson-Crick base pairing of the incoming NTP with the template-RNA (t-RNA) base closes the active site. This conformational change promotes nucleophilic attack of the p-RNA 3’-OH on the NTP α-phosphate, releasing a pyrophosphate in a S_N_2 condensation reaction ^11^. Available SARS-CoV-2 RTC structures are dominated by apo structures with an empty NTP active site lacking catalytic metals and with an open active-site conformation, hindering structure-based drug design ^14,15,16,17^. Only two studies document the RTC in the presence of substrates: the antiviral inhibitors favipiravir triphosphate and the activated form of AT-527, AT-9010, but neither investigation yielded views of a catalytically competent RTC due to unproductive binding modes and open conformations of the active sites ^18,19^. Therefore, to gain mechanistic insights into RTC recognition of its substrates during RNA synthesis, we determined five cryo-electron microscopy (cryo-EM) structures, capturing the RTC with each of the four natural NTPs as well as with RDV-TP. The five cryo-EM maps range in nominal resolutions between ∼2.6-3.3 Å, with the RdRp active sites resolved locally to ∼2.2-2.9 Å (Extended Data Fig. 1-5, Extended Data Table 1) ^20^, enabling near-atomic resolution insights into the mechanism of NTP recognition and RDV-TP selectivity.

**Table 1.** Cryo-EM data collection, refinement and validation statistics.

|  | **S1_RDV-TP** | **S2_ATP** | **S3_UTP** | **S4_GTP** | **S5_CTP** |
| --- | --- | --- | --- | --- | --- |
|  | EMDB-26639 | EMD-26641 | EMD-26642 | EMDB-26645 | EMDB-26646 |
|  | PDB 7UO4 | PDB 7UO7 | PDB 7UO9 | PDB 7UOB | PDB 7UOE |
| **Data collection and processing** |  |  |  |  |  |
| **Magnification** | 81,000 | 81,000 | 81,000 | 64,000 | 130,000 |
| **Voltage (kV)** | 300 | 300 | 300 | 300 | 300 |
| **Electron exposure (e<sup>-</sup>/Å<sup>2</sup>)** | 54.58 | 51.89 | 53.34 | 51.44 | 56.767 |
| **Defocus range (µm)** | -0.8 µm to -2.5 µm | -0.8 µm to -2.5 µm | -0.8 µm to -2.5 µm | -0.8 µm to -3.0 µm | -0.8 µm to -3.0 µm |
| **Pixel size (Å)** | 1.065 | 1.076 | 1.069 | 1.08 | 0.515 |
| **Symmetry imposed** | C1 | C1 | C1 | C1 | C1 |
| **Initial particle images (no.)** | 8,017,151 | 5,363,858 | 14,149,078 | 2,419,929 | 1,986,527 |
| **Final particle images (no.)** | 613,848 | 330,442 | 719,899 | 456,629 | 171,107 |
| **Map resolution (Å)**<br>**FSC threshold 0.143** | 3.38 | 3.09 | 3.13 | 2.68 | 2.67 |
| **Map resolution range (Å)** | 2.91 - 6.68 | 2.82 - 7.53 | 2.74 - 6.81 | 2.29 - 7.82 | 2.34 - 8.24 |
| **Refinement** |  |  |  |  |  |
| **Initial models used (PDB code)** | 7RE1 | 7RE1 | 7RE1 | 7RE1 | 7RE1 |
| **Model resolution (Å)**<br>**FSC threshold 0.5** | 3.5 | 3.4 | 3.2 | 2.8 | 2.9 |
| **Map sharpening B factor (Å<sup>2</sup>)** | 144.9 | 100.7 | 141.4 | 106.7 | 90.6 |
| **Model composition** |  |  |  |  |  |
| **Non-hydrogen atoms** | 12,470 | 12,262 | 12,560 | 12,718 | 12,705 |
| **Protein residues** | 1,374 | 1,371 | 1,375 | 1,375 | 1,376 |
| **Nucleic acid residues (RNA)** | 70 | 66 | 67 | 69 | 70 |
| **Ligands** | 4 | 4 | 4 | 7 | 5 |
| **B factors (Å<sup>2</sup>)** |  |  |  |  |  |
| **Protein** | 23.34 | 74.21 | 37.93 | 16.96 | 57.92 |
| **Nucleic acids** | 116.61 | 153.2 | 117.39 | 82.17 | 143.63 |
| **Ligand** | 16.71 | 58.48 | 25.68 | 13.75 | 48.02 |
| **R.m.s. deviations** |  |  |  |  |  |
| **Bond lengths (Å)** | 0.004 | 0.005 | 0.005 | 0.005 | 0.004 |
| **Bond angles (°)** | 0.503 | 0.643 | 0.508 | 0.605 | 516 |
| **Validation** |  |  |  |  |  |
| **MolProbity score** | 1.93 | 1.8 | 1.65 | 1.45 | 1.44 |
| **Clashscore** | 7.28 | 8.32 | 4.94 | 3.27 | 4.95 |
| **Poor rotamers (%)** | 1.27 | 0.76 | 0.25 | 1.52 | 1.18 |
| **Ramachandran plot** |  |  |  |  |  |
| **Favored (%)** | 92.83 | 95.01 | 94.15 | 96.78 | 97.37 |
| **Allowed (%)** | 7.17 | 4.99 | 5.85 | 3 | 2.41 |
| **Disallowed (%)** | 0 | 0 | 0 | 0.22 | 0.22 |

## Trapping SARS-CoV-2 RTC pre-incorporation complexes

Guided by previous mechanistic work on nucleic acid polymerases, we investigated a battery of chemical strategies to block incorporation after substrate binding. These approaches included using nucleotide diphosphate (NDP) substrates ^13^, alternative metal co-factors (such as Ca^2+^) ^21^, α-β non-hydrolysable nucleotide analogues ^22^, and 3’-deoxy p-RNA scaffolds. Although the use of NDP substrates trapped the hepatitis C virus (HCV) RdRp recognition complex ^13^, the SARS-CoV-2 RTC efficiently incorporated GDP, less efficiently ADP, and to a minor extent RDV-diphosphate (RDV-DP) (Extended Data Fig. 6, see also ^23^). The ability of the SARS-CoV-2 RTC to incorporate NDPs reflects the relative promiscuity of the RdRp active site, which also retained RNA synthesis activity in the presence of both Ca^2+^ and α-β-imino analogues (Extended Data Fig. 6e). However, RTC synthesis activity was fully ablated with the incorporation of a 3’-deoxy nucleotide into the p-RNA, leading us to utilize this approach to visualize the RTC pre-incorporation complexes (Extended Data Fig. 6g).

To validate the effectiveness of using 3’-deoxy RNA scaffolds, we used native mass spectrometry (nMS) to probe for the extension of the primer by one base or the formation of a stalled ternary complex (Fig 1b, c, Extended Data Fig. 7) in the presence of various NTPs. In the event of nucleotide incorporation, we expected a mass-shift corresponding to NMP addition, whereas non-covalently bound (unincorporated) substrates would either dissociate or yield a mass-shift corresponding to the NTP and bound metal ions. For the control scaffolds containing a 3’-OH, near complete p-RNA extension by a single NMP occurred (Fig. 1b). In the presence of p-RNA strands lacking a 3’-OH, no incorporation of the next nucleotide (ATP, RDV-TP, or GTP) was observed, confirming the suitability of this strategy for trapping pre-incorporation complexes (Fig. 1c). In the pre-incorporation complexes, we observed peaks corresponding to the RTC with the incoming Mg-NTP (ATP, RDV-TP, or GTP) with the relative peak intensity in the order of GTP > RDV-TP > ATP (Fig. 1c).

## Structural basis of NTP recognition

Having identified a strategy to capture substrate-bound RTCs stalled in a pre-incorporation state, we prepared cryo-EM samples of RTCs with each of the natural nucleotides, ATP, CTP, GTP, UTP, or with RDV-TP (Extended Data Fig. 1-5). The resulting structures yielded the first views of the RTC poised for catalysis. Two of the structures, the RTC with CTP (+CTP) or GTP (+GTP), yielded nominal resolutions better than 3 Å (2.7 Å), with local resolution around the active site estimated to be ∼2.2 Å ^20^, enabling near-atomic resolution insights into substrate recognition.

Density features in the +CTP and +GTP cryo-EM maps are consistent with the positions of divalent metal ions observed in many previous nucleic acid polymerase structures ^11-13, 24^. Based on the presence of 5 mM MgCl_2_ in the cryo-EM buffer and the observed octahedral coordination geometries, we assigned these densities as Mg^2+^_A_ and Mg^2+^_B_ (Fig. 2b, Supplementary Video 1), consistent with the hypothesized universal two-metal ion mechanism for nucleic acid polymerases ^11^. The density peak for Mg^2+^_A_ in the +CTP and +GTP structures is weaker than that for Mg^2+^_B_, and density for Mg^2+^_A_ in the +ATP, +UTP, and +RDV-TP structures is completely absent. Despite the absence of Mg^2+^_A_ in the +ATP, +UTP, and +RDV-TP structures, most details of the active-site configurations were indistinguishable from the +CTP and +GTP structures.

**Fig. 2.**
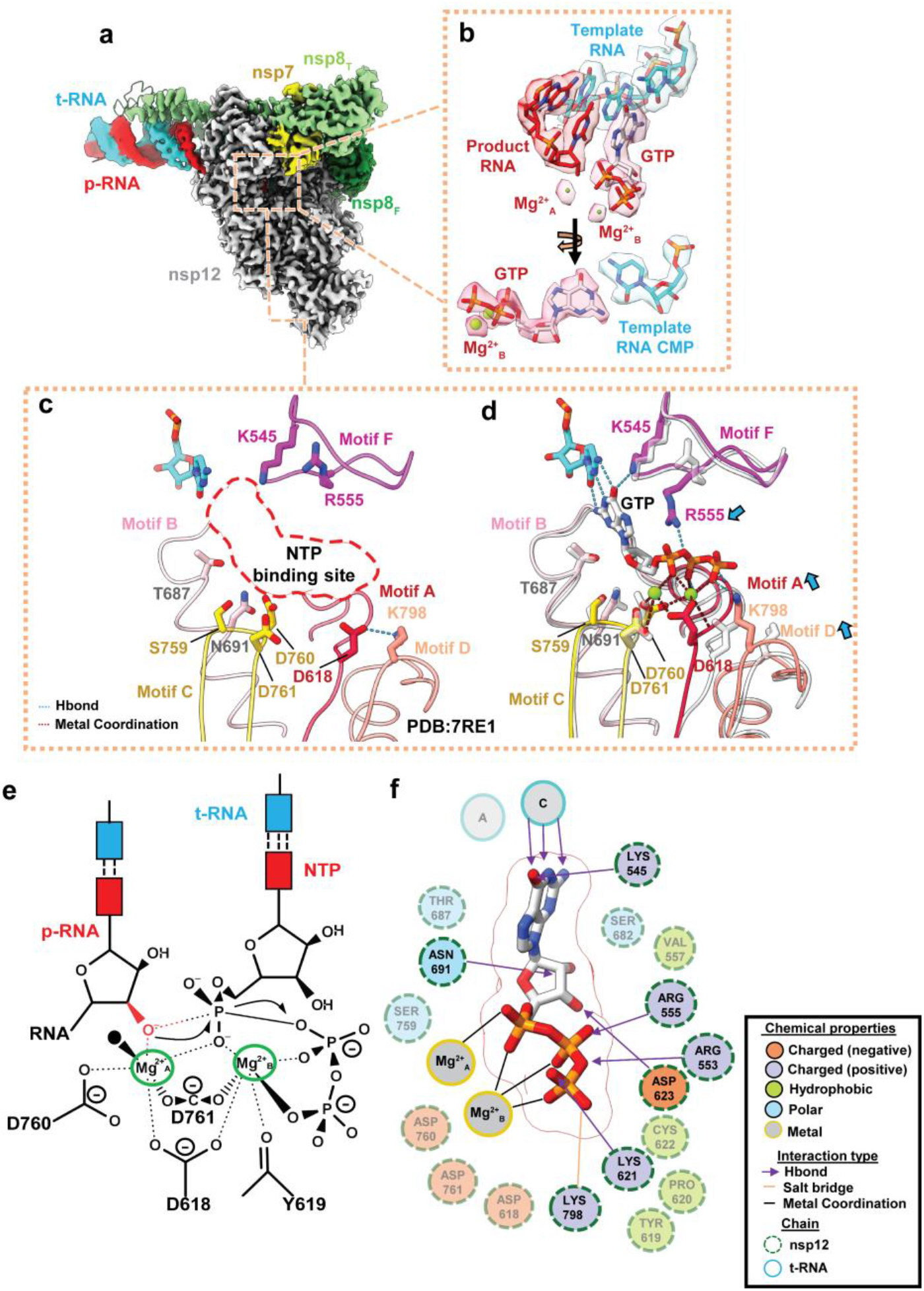
Structural basis of nucleotide recognition. **(a)** Cryo-EM density of the 2.7 Å nominal resolution +GTP structure (S4_GTP), colored according to the fitted model chains **(b)** Cryo-EM density of the bound incoming GTP, two associated metal ions, and the nearby bases of the template-RNA (t-RNA) and product-RNA (p-RNA) strands. **(c)** Close-up of the active site of the apo complex PDB 7RE1 ^(17)^, illustrating the arrangement of the conserved RdRp active site motifs A-D and F (crimson, hot pink, gold, salmon, magenta, respectively) and of key residues when the NTP binding site (dashed red line) is empty. **(d)** Comparison of the S4_GTP and apo structure (faded grey) active sites, highlighting observed motif/residue rearrangements (blue arrows) on NTP binding. Movements in motifs A and D close the active site. **(e)** Schematic depicting nucleotide addition, in the presence of an intact 3’-OH, based on disposition of magnesiums and NTP in the S4_GTP structure **(f)** 2D schematic of the suite of interactions involved in NTP binding based on distances in the S4_GTP structure. Residues interacting indirectly with the NTP are faded for clarity.

A consistent feature observed across all five structures is the closure of the active site around the incoming NTP, mediated by a rotation of the RdRp motif A towards the NTP substrate (Fig. 2c, d). Closure of motif A stabilizes substrate binding by: (I) enabling the backbone carbonyl of Y619 to coordinate Mg^2+^_B_ and the catalytic residue D618 to coordinate both Mg^2+^_A_ and Mg^2+^_B_ (Fig. 2e), (II) promoting the formation of a hydrogen-bonding (H-bonding) network through D623 that enables the recognition of the substrate ribose 3’-OH by S682 and N691 (motif B), (III) enabling weak H-bonding interactions between the β- and γ-phosphates and motif A residues K621 or C622 (Fig. 2f, Extended Data Fig. 9f); and (IV) disrupting the polar D618-K798 interaction in the apo-RTC^14, 15, 17^, repositioning D618 for metal coordination, and K798 to interact with the NTP γ-phosphate (Fig. 2d).

The detailed interactions of nsp12 residues with the four natural NTP substrates were essentially identical except for conserved basic residues of motif F. K545 (motif F) displays selective interactions with the incoming NTPs, forming a H-bond with carbonyl oxygens of UTP (C4) and GTP (C6), interactions that are precluded by the amino group at these positions in the ATP, RDV-TP, or CTP bases (Extended Data Fig. 9). The cryo-EM densities also suggest that the side chains of K551 and R555 are dynamic, accessing multiple conformations in each structure. K551 primarily interacts with the γ-phosphate of the CTP and RDV-TP substrates, but weak density indicates disorder in the other structures. In the +ATP and +RDV-TP structures, R555 predominantly forms a pi-pi stacking interaction with the NTP substrate base. This differs from the predominant conformation in the +CTP, +GTP, and +UTP structures where the side chain of R555 forms a H-bond with the β-phosphate oxygens of the NTP substrate (Fig. 3, Extended data Fig. 9). In either state, R555 buttresses the incoming nucleotide, arranging the active site for catalysis. We speculate that the observed R555 stacking interaction may first promote the formation of a canonical Watson-Crick base pair by stabilizing the nucleobase opposite the template base, then reorient to interact with the nucleotide β-phosphate to promote subsequent catalysis by reinforcing the correct geometry of the pyrophosphate leaving group, roles that may be conserved across RdRps such as HCV RdRp ^25^.

**Fig. 3.**
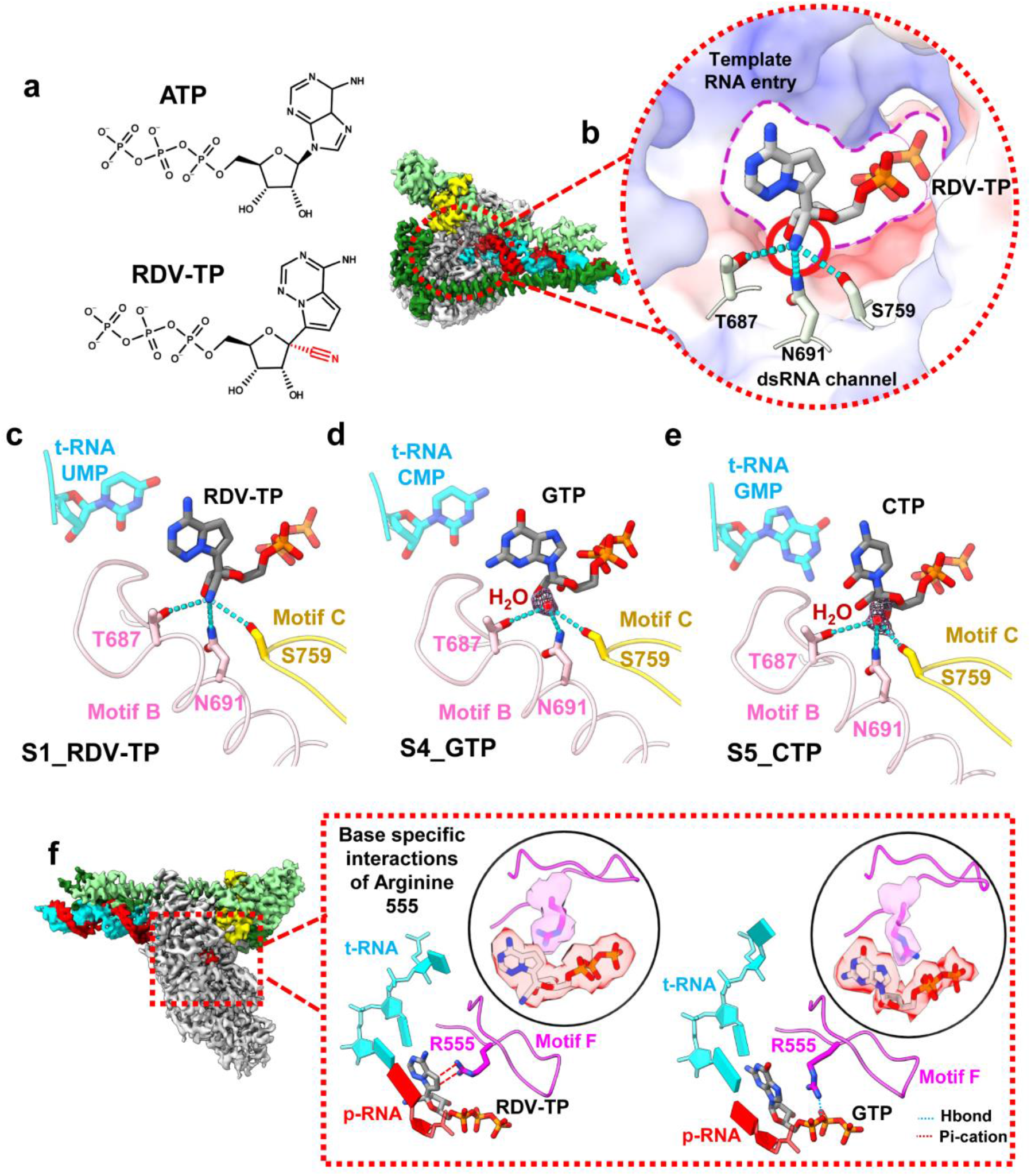
Molecular basis of Remdesivir’s incorporation selectivity. **(a)** Chemical structures of ATP and RDV-TP, highlighting the position of the RDV-TP 1’ cyano group. **(b)** Cryo-EM densities of the S2_RDV-TP structure, colored according to fitted model chains. Zoom-in on the bound RDV-TP illustrates the RDV-TP 1’ cyano group is accommodated in a hydrophilic pocket formed by motif B and C residues. Protein surface is colored according to electrostatics. **(c-e)** Comparison of the active sites of the S1_RDV-TP **(c)**, S4_GTP **(d)**, S5_CTP **(e)** structures reveals that the RdRp cyano pocket can also bind a water molecule (map density around the water shown in mesh) which needs to be displaced for RDV-TP binding **(f)** A comparison of the S2_RDV-TP and S4_GTP structures reveals two predominant rotamers of R555 which mediate either a pi-pi stacking (red dotted lines) or a H-bond (blue dotted line) interaction with the incoming NTP.

The structural observation of SARS-CoV-2 RdRp motif A closure resembles a similar motion of motif A observed in substrate-bound RdRp complexes of polio virus and HCV ^12, 13^, illustrating the universal nature of the RdRp palm closure step during NTP recognition. Although the palm-mediated active site closure is well-documented, our results indicate that the structural plasticity of positively charged residues in motif F (K545, K551, R553, and R555) in the fingers domain helps orient the NTP substrate for catalysis through interactions with both the nucleobase and triphosphate moieties. We note that these residues are invariant across the major coronavirus clades and GISAID database of patient SARS-CoV-2 sequences (Extended Data Fig. 8), reflecting the near immutable nature of the coronavirus RdRp active site. Previously, we noted that the same residues line the NTP entry channel following nsp13-mediated p-RNA backtracking ^26^, illustrating the pleiotropic role that motif F plays during RNA synthesis.

## Structural basis of RDV-TP selectivity

RDV-TP has been characterized in biochemical studies to possess an almost threefold greater selectivity for incorporation into elongating RNA compared to ATP ^3, 4^, a property which is thought to improve its ability to inhibit RNA synthesis. In our structures, both nucleotides are observed base-paired to the cognate t-RNA U; the two nucleotides superimpose with a root-mean-square-deviation (RMSD) of 0.34 Å over 32 common atoms. The key difference between RDV-TP and ATP is the 1’-cyano moiety on the RDV-TP ribose (Fig. 3a), which juts into a hydrophilic pocket formed by residues T687, N691 (motif B), and S759 (motif C) (Fig. 3c), creating a network of polar interactions (Fig. 3c). In the highest resolution cryo-EM maps (+CTP and +GTP; Extended Data Table 1), we observed a stably bound water molecule occupying the hydrophilic pocket in the absence of the RDV-TP cyano-group (compare Fig. 3c to 3d, e), suggesting that the enhanced affinity (lower *K*_*m*_) for RDV-TP over ATP ^3, 4^ may be attributed to both an entropic effect through release of the bound water and an enthalpic effect through formation of new polar interactions between RDV-TP and the surrounding polar residues. A recent study reports that SARS-CoV-2 acquired phenotypic resistance to RDV through an nsp12-S759A substitution following serial passage in cell culture ^27^, indicating that binding of the RDV-TP 1’-cyano group in this hydrophilic pocket is an important facet of RDV susceptibility.

## Structural insights into RNA capping

Apart from the RdRp active site, the RTC possesses an additional catalytic domain known as the NiRAN (Nidovirus RdRp-associated nucleotidyltransferase), which lies N-terminal of the RdRp ^5, 14, 28^. Although the NiRAN domain is essential for viral propagation, its functions during the viral life cycle remain enigmatic. Recent studies have suggested that the NiRAN functions in the viral RNA capping pathway in conjunction with an additional viral protein, nsp9 ^5, 29^. Others have postulated a role for the NiRAN domain in protein-mediated priming of RNA synthesis ^30^. In previous structural studies, ADP ^14^ or GDP ^29^ have been observed bound in the NiRAN active site in a ‘base-out’ pose, where the nucleotide base points out of the active site pocket and makes few, if any, protein contacts. ADP has not been shown to serve as a substrate for the NiRAN enzymatic activity, and the GDP was bound along with an N-terminally modified (and resultingly inactive) nsp9 ^29^, so the relevance of these nucleotide poses is unclear. In another study, the diphosphate form of the GTP analog AT-9010 is seen bound to the NiRAN in a ‘base-in’ pose ^19^, with the phosphates occupying the same positions as the phosphates in the ‘base-out’ pose but flipped, and with the guanine base bound in a tight pocket internal to the NiRAN (Extended Data Fig. 10).

In our cryo-EM structures of the RTC with each of the natural NTPs and RDV-TP (Extended Data Table 1), only GTP was observed stably and specifically bound in the NiRAN active site. The cryo-EM density supports the presence of GTP (and not GDP) in the NiRAN (Fig. 4). The GTP is bound in the ‘base-in’ pose similar to the diphosphate form of AT-9010 ^19^, with the base buried in an apparently guanine-specific pocket that extends into the core of the NiRAN fold, enabling it to make key contacts with hydrophilic residues that line the pocket interior. A central element to this recognition is interactions mediated by R55 and Y217, two conserved residues across the α- and β-coronavirus clades, which provide base specificity for guanosine (Fig. 4 b, d). The apo-NiRAN pocket is sterically incompatible with the bound guanine base; GTP association is mediated by an induced fit mechanism that involves expansion of the active-site pocket to accommodate the guanosine base (Fig. 4e).

**Fig. 4.**
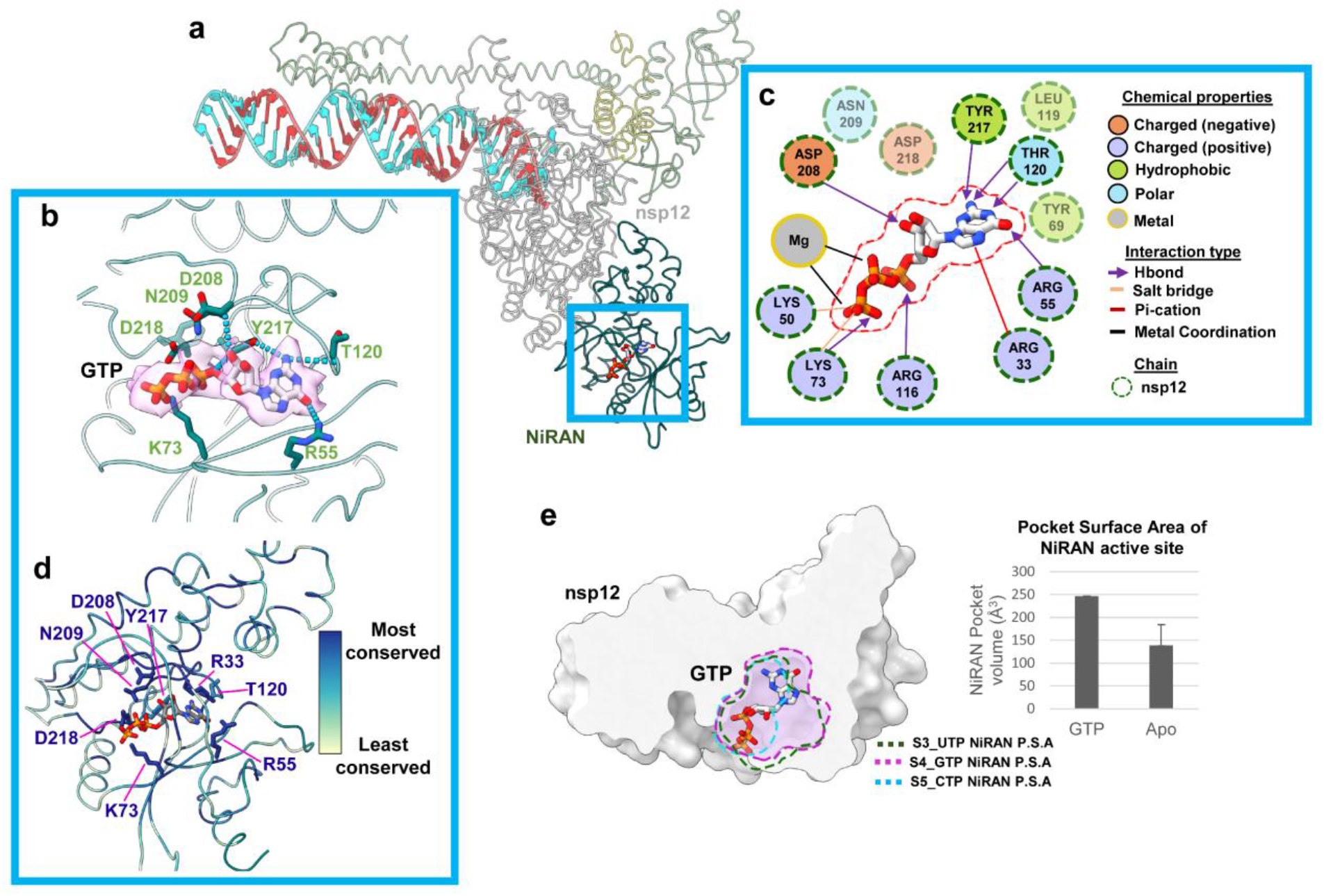
NiRAN specific recognition of GTP. **(a)** View of the NiRAN domain of the RTC, bound to GTP, which lies at the amino-terminal end of nsp12. **(b)** GTP is selectively recognized in the NiRAN pocket by a series of hydrogen bonding and electrostatic interactions in which interacting residues are shown as sticks. **(c)** 2D schematic illustrating the NiRAN-GTP interactions. **(d)** Residues that mediate GTP recognition are shown as sticks and colored according to their conservation across the α & β coronavirus clades. **(e)** Binding of GTP in the NiRAN is mediated through an induced fit that widens the pocket for insertion of the guanine base. Cross-sections (dashed lines) of the pocket surface area (P.S.A) of the S4_GTP (GTP bound), S3_UTP and S5_CTP (apo-NiRAN) are shown overlaid on a clipped surface view of the nsp12 NiRAN. The NiRAN pocket volume was measured using the Schrodinger Sitemap tool.

A recent preprint proposes that the NiRAN mediates two successive steps in the viral RNA capping pathway that initially entails transfer of the nascent RNA 5’-end to the amino terminus of nsp9, forming a covalent RNA-protein intermediate in a process termed RNAylation ^6^. A successive polyribonucleotidyltransferase (PRNTase) reaction utilizes the RNAylated nsp9 as a substrate and transfers the bound RNA to GDP/GTP to produce the cap structure, GpppN-RNA. The PRNTase activity strictly utilizes GDP or GTP. Although the authors report slightly higher PRNTase activity when GDP is the substrate, GTP may be the more likely physiological substrate given that concentrations of GTP are much higher than GDP in the cellular milieu ^31^. Our results indicate that the NiRAN uniquely recognizes GTP (but not other NTPs), explaining the base specificity observed for the PRNTase activity.

## Discussion

Antiviral nucleotide analogues are highly effective and versatile therapeutics that can be adapted to treat novel coronaviral borne diseases as they emerge. Their suitability for repurposing is exemplified by the use of RDV and molnupiravir for the treatment of COVID-19, two analogues that were initially developed as therapeutics against Ebola virus ^32^ and influenza virus ^33^. Repurposing efforts rely on the conserved active site of the RdRp, which has not been observed to mutate as readily as other COVID-19 therapeutic targets such as the Spike protein (Extended Data Fig. 8). While active site residues are highly conserved across RdRp-encoding viruses, subtle differences can alter the incorporation selectivity of a nucleotide analogue more than 100-fold ^4^ highlighting the importance of mechanistic investigations of RdRps across the viral realm.

To probe the mechanism of nucleotide recognition by the SARS-CoV-2 RdRp, we determined structures of stalled RdRp complexes containing an RNA primer-template, incoming nucleotide substrates or the antiviral RDV-TP, and catalytic metal ions. Structural views of the RTC with each of the respective nucleotides illustrate how the RdRp structurally adapts to proper Watson-Crick base pairing geometry, closing in around the t-RNA/NTP base pair to facilitate catalytic Mg^2+^ coordination and in-line attack of the p-RNA 3’-OH on the NTP α-phosphate.

Through visualizing these structures, we ascertained that the enhanced selectivity for RDV, which primarily materializes biochemically as an effect on *K*_*m*_, is mediated by the accommodation of its 1’-cyano group in a conserved hydrophilic pocket near the RdRp active site (Fig. 3). This same RdRp active site pocket harbors an ordered water molecule in the +CTP and +GTP structures (Fig. 3); the bound water is likely present in the +ATP and +UTP complexes as well but not visualized due to limited resolution (Extended Data Figs. 2, 3). Mutations that alter the hydrophilicity of this pocket give rise to RDV resistance ^27^ and are naturally found in viral families that exhibit reduced sensitivity to RDV ^4^. Critically, structural insights into NTP recognition could be exploited in the design of nucleotide analogues as part of structure-based drug design programs that aim to target the coronavirus RdRp. Our studies also shed light on the presence of a guanosine-specific pocket in the essential NiRAN domain, supporting proposals that the NiRAN is involved in the production of capped mRNAs as well as further detailing this pocket as a therapeutic target.

## METHODS

No statistical methods were used to predetermine sample size. The experiments were not randomized, and the investigators were not blinded to allocation during experiments and outcome assessment.

### Protein expression and purification

*SARS-CoV-2 nsp12* was expressed as previously described ^34^ and purified as follows. Briefly, a pQE-30/pcl-ts ind+ plasmid containing a His_6_-SUMO SARS-CoV-2 nsp12 and untagged nsp7 & 8 (Addgene #160540) was transformed into *E. coli* BL21 cells (Agilent). Cells were grown and protein expression was induced by the addition of 0.2 mM isopropyl β-d-1-thiogalactopyranoside (IPTG), 10 ng/mL tetracycline and 50 ug/mL nalidixic acid. Cells were collected and lysed in a French press (Avestin). The lysate was cleared by centrifugation and purified on a HiTrap Heparin HP column (Cytiva). The fractions containing nsp12 were loaded onto a HisTrap HP column (Cytiva) for further purification. Eluted nsp12 was dialyzed, cleaved with His_6_-Ulp1 SUMO protease, and passed through a HisTrap HP column to remove the SUMO protease. Flow-through was collected, concentrated by centrifugal filtration (Amicon), and loaded on a Superdex 200 Hiload 16/600 (Cytiva). Glycerol was added to the purified nsp12, aliquoted, flash-frozen with liquid N_2_, and stored at -80°C.

*SARS-CoV-2 nsp7/8* was expressed and purified as described ^14^. Briefly, the pCDFDuet-1 plasmid containing His_6_ SARS-CoV-2 nsp7/8 (Addgene #159092) was transformed into *E. coli* BL21 (DE3). Cells were grown and protein expression was induced by the addition of IPTG. Cells were collected and lysed in a French press (Avestin). The lysate was cleared by centrifugation and purified on a HisTrap HP column (Cytiva). Eluted nsp7/8 was dialyzed, cleaved with His_6_-Prescission Protease to cleave His_6_ tag, and then passed through a HisTrap HP column to remove the protease (Cytiva). Flow-through was collected, concentrated by centrifugal filtration (Amicon), and loaded onto a Superdex 75 Hiload 16/600 (Cytiva). Glycerol was added to the purified nsp7/8, aliquoted, flash-frozen with liquid N_2_, and stored at -80°C.

### Preparation of SARS-CoV-2 replication/transcription complex (RTC) for Cryo-EM

Cryo-EM samples of SARS-CoV-2 RTC were prepared as previously described ^14, 17^. Briefly, purified nsp12 and nsp7/8 were concentrated, mixed in a 1:3 molar ratio, and incubated for 20 min at 22°C. An annealed RNA scaffold (Horizon Discovery, Ltd.) was added to the nsp7/8/12 complex and incubated for 10 min at 30°C. Sample was buffer exchanged into cryo-EM buffer [20 mM HEPES pH 8.0, 100 mM K-Acetate, 5 mM MgCl_2_, 2 mM DTT] and further incubated for 20 min at 30°C. The sample was purified over a Superose 6 Increase 10/300 GL column (Cytiva) in cryo-EM buffer. The peak corresponding to nsp7/8/12/RNA complex was pooled and concentrated by centrifugal filtration (Amicon).

### Cryo-EM grid preparation

Prior to grid freezing, beta-octyl glucoside (β-OG) was added to the sample (0.07 % w/v final). The final buffer condition for the cryo-EM sample was 20 mM HEPES pH 8.0, 100 mM K-Acetate, 5 mM MgCl_2_, 2 mM DTT, 0.07% (w/v) β-OG with 300 μM final of the respective NTP added immediately before freezing. C-flat holey carbon grids (CF-1.2/1.3-4Au, EMS) were glow-discharged for 20 seconds prior to the application of 3.5 μL of sample. Using a Vitrobot Mark IV (Thermo Fisher Scientific), grids were blotted and plunge-frozen into liquid ethane with 95% chamber humidity at 4°C.

### Cryo-EM data acquisition and processing

Structural biology software was accessed through the SBGrid consortium ^35^. The following pertains to each of the respective datasets:

### S1_RDV-TP

Grids were imaged using a 300 kV Titan Krios (Thermo Fisher Scientific) equipped with a GIF BioQuantum and K3 camera (Gatan). Images were recorded with Leginon ^36^ with a pixel size of 1.065 Å/px (micrograph dimension of 5760 × 4092 px) over a defocus range of −0.8 μm to −2.5 μm with a 20 eV energy filter slit. Movies were recorded in “counting mode” (native K3 camera binning 2) with ∼25 e-/px/s in dose-fractionation mode with subframes of 50 ms over a 2.5 s exposure (50 frames) to give a total dose of ∼54 e-/Å^2^. Dose-fractionated movies were gain-normalized, drift-corrected, summed, and dose-weighted using MotionCor2 ^37^. The contrast transfer function (CTF) was estimated for each summed image using the Patch CTF module in cryoSPARC v3.1.0 ^38^. Particles were picked and extracted from the dose-weighted images with box size of 256 px using cryoSPARC Blob Picker and Particle Extraction. The entire dataset consisted of 26,130 motion-corrected images with 8,017,151 particles. Particles were sorted using three rounds of cryoSPARC 2D classification (N=50, where N equals the number of classes), resulting in 2,473,065 curated particles. Initial models (denoted as monomer & dimer) were generated using cryoSPARC *Ab initio* Reconstruction (N=3) on a subset of the particles. Particles were further curated using these initial models as 3D templates for iterative cryoSPARC Heterogeneous Refinement (N=4), resulting in 314,848 particles in the monomer class (cyan map, Extended Data Fig. 1) and 264,453 particles in the dimer class (pink map, Extended Data Fig. 1). Curated particles in the monomer and dimer classes were re-extracted to a box size of 384 and input to cryoSPARC Homogenous and Non-uniform refinements ^39^. Particles within each class were further processed through two rounds of RELION 3.1 Bayesian Polishing ^40^. Polished particles were refined using cryoSPARC Local and Global CTF Refinement in combination with cryoSPARC Non-uniform Refinement, resulting in structures with the following particle counts and nominal resolutions: monomer RTC (251,160 particles; 3.54 Å) & dimer RTC (249,468 particles; 3.87 Å). To facilitate model building of the RTC, particles from the dimer RTC class underwent masked particle subtraction in which both protomers were masked, subtracted from the full map, and combined with the original monomer RTC to yield a structure of the RTC at a nominal resolution of 3.38 Å from 613,848 particles (Extended Data Fig. 1e). Local resolution calculations were generated using blocres and blocfilt from the Bsoft package ^20^ (Extended Data Fig. 1c). The angular distribution of particle orientations (Extended Data Fig. 1b) and directional resolution through the 3DFSC package ^41^ (Extended Data Fig. 1d) were calculated for the final class.

### S2_ATP

Grids were imaged using a CS corrected 300 kV Titan Krios (Thermo Fisher Scientific) equipped with a GIF BioQuantum and K3 camera (Gatan). Images were recorded with Leginon ^36^ with a pixel size of 1.076 Å/px (micrograph dimension of 5760 × 4092 px) over a defocus range of −0.8 μm to −2.5 μm with a 20 eV energy filter slit. Movies were recorded in “counting mode” (native K3 camera binning 2) with ∼25 e-/px/s in dose-fractionation mode with subframes of 50 ms over a 2.5 s exposure (50 frames) to give a total dose of ∼52 e-/Å^2^. Dose-fractionated movies were gain-normalized, drift-corrected, summed, and dose-weighted using MotionCor2 ^37^. The CTF was estimated for each summed image using the Patch CTF module in cryoSPARC v3.1.0 ^38^. Particles were picked and extracted from the dose-weighted images with box size of 256 px using cryoSPARC Blob Picker and Particle Extraction. The entire dataset consisted of 11,657 motion-corrected images with 5,364,858 particles. Particles were sorted using three rounds of cryoSPARC 2D classification (N=50), resulting in 2,255,856 curated particles. Initial models (denoted as monomer & dimer) were generated using cryoSPARC *Ab initio* Reconstruction (N=4) on a subset of the particles. Particles were further curated using these initial models as 3D templates for iterative cryoSPARC Heterogeneous Refinement (N=4), resulting in 102,529 particles in the monomer class (cyan map, Extended Data Fig. 2) and 318,672 particles in the dimer class (pink map, Extended Data Fig. 2). Curated particles in the monomer and dimer classes were re-extracted to a box size of 384 and input to cryoSPARC Homogenous and Non-uniform refinements ^39^. Particles within each class were further processed through two rounds of RELION 3.1 Bayesian Polishing ^40^. Polished particles were refined using cryoSPARC Local and Global CTF Refinement in combination with cryoSPARC Non-uniform Refinement, resulting in structures with the following particle counts and nominal resolutions: monomer RTC (96,868 particles; 3.71 Å) & dimer RTC (299,965 particles; 3.50 Å). To facilitate model building of the RTC, particles from the dimer RTC class underwent masked particle subtraction in which both protomers were masked, subtracted from the full map, and combined with the original monomer RTC to yield a structure of the RTC at a nominal resolution of 3.09 Å from 330,442 particles (Extended Data Fig. 2e). This combined RTC class was further refined with masked cryoSPARC Local Refinement using masks around the RdRp and NiRAN active sites. Locally refined maps were combined into a RTC composite map using PHENIX ‘Combine Focused Maps’ to aid model building ^42^. Local resolution calculations were generated using blocres and blocfilt from the Bsoft package ^20^ (Extended Data Fig. 2c). The angular distribution of particle orientations (Extended Data Fig. 2b) and directional resolution through the 3DFSC package ^41^ (Extended Data Fig. 2d) were calculated for the final class.

### S3_UTP

Grids were imaged using a 300 kV Titan Krios (Thermo Fisher Scientific) equipped with a GIF BioQuantum and K3 camera (Gatan). Images were recorded with Leginon ^36^ with a pixel size of 1.065 Å/px (micrograph dimension of 5760 × 4092 px) over a defocus range of −0.8 μm to −2.5 μm with a 20 eV energy filter slit. Movies were recorded in “counting mode” (native K3 camera binning 2) with ∼25 e-/px/s in dose-fractionation mode with subframes of 50 ms over a 2.5 s exposure (50 frames) to give a total dose of ∼53 e-/Å^2^. Dose-fractionated movies were gain-normalized, drift-corrected, summed, and dose-weighted using MotionCor2 ^37^. The CTF was estimated for each summed image using the Patch CTF module in cryoSPARC v3.1.0 ^38^. Particles were picked and extracted from the dose-weighted images with box size of 256 px using cryoSPARC Blob Picker and Particle Extraction. The entire dataset consisted of 30,850 motion-corrected images with 14,149,078 particles. Particles were sorted using three rounds of cryoSPARC 2D classification (N=50), resulting in 3,297,109 curated particles. Initial models (denoted as monomer & dimer) were generated using cryoSPARC *Ab initio* Reconstruction (N=4) on a subset of the particles. Particles were further curated using these initial models as 3D templates for iterative cryoSPARC Heterogeneous Refinement (N=6), resulting in 648,814 particles in the monomer class (cyan map, Extended Data Fig. 3) and 773,911 particles in the dimer class (pink map, Extended Data Fig. 3). Curated particles in the monomer and dimer classes were re-extracted to a box size of 384 and input to cryoSPARC Homogenous and Non-uniform refinements ^39^. Particles within each class were further processed through two rounds of RELION 3.1 Bayesian Polishing ^40^. Polished particles were refined using cryoSPARC Local and Global CTF Refinement in combination with cryoSPARC Non-uniform Refinement, resulting in structures with the following particle counts and nominal resolutions: monomer RTC (614,648 particles; 3.34 Å) & dimer RTC (730,009 particles; 3.42 Å). To facilitate model building of the RTC, particles from the dimer RTC class underwent masked particle subtraction in which both protomers were masked, subtracted from the full map, and combined with the original monomer RTC to yield a structure of the RTC at a nominal resolution of 3.13 Å from 719,889 particles (Extended Data Fig. 3e). This combined RTC class was further refined with masked cryoSPARC Local Refinement using masks around the RdRp and NiRAN active sites. Locally refined maps were combined into a RTC composite map using PHENIX ‘Combine Focused Maps’ to aid model building ^42^. Local resolution calculations were generated using blocres and blocfilt from the Bsoft package ^20^ (Extended Data Fig. 3c). The angular distribution of particle orientations (Extended Data Fig. 3b) and directional resolution through the 3DFSC package ^41^ (Extended Data Fig. 3d) were calculated for the final class.

### S4_GTP

Grids were imaged using a CS corrected 300 kV Titan Krios (Thermo Fisher Scientific) equipped with a GIF BioQuantum and K3 camera (Gatan). Images were recorded with SerialEM ^43^ with a pixel size of 1.08 Å/px (micrograph dimension of 5760 × 4092 px) over a defocus range of −0.8 μm to −3.0 μm with a 20 eV energy filter slit. Movies were recorded in “counting mode” (native K3 camera binning 2) with ∼25 e-/px/s in dose-fractionation mode with subframes of 50 ms over a 2.5 s exposure (50 frames) to give a total dose of ∼51 e-/Å^2^. Dose-fractionated movies were gain-normalized, drift-corrected, summed, and dose-weighted using MotionCor2 ^37^. The CTF was estimated for each summed image using the Patch CTF module in cryoSPARC v3.1.0 ^38^. Particles were picked and extracted from the dose-weighted images with box size of 256 px using cryoSPARC Blob Picker and Particle Extraction. The entire dataset consisted of 4,527 motion-corrected images with 2,419,929 particles. Particles were sorted using three rounds of cryoSPARC 2D classification (N=50), resulting in 941,507 curated particles. An initial model of the monomer RTC was generated using cryoSPARC *Ab initio* Reconstruction (N=3) on a subset of the particles. Particles were further curated using this initial model as a 3D template for iterative cryoSPARC Heterogeneous Refinement (N=6), resulting in 484,682 particles in the resultant monomer class (cyan map, Extended Data Fig. 4). Curated particles were re-extracted to a box size of 384 and input to cryoSPARC Homogenous and Non-uniform refinements ^39^. Particles were further processed through two rounds of RELION 3.1 Bayesian Polishing ^40^. Polished particles were refined using cryoSPARC Local and Global CTF Refinement in combination with cryoSPARC Non-uniform Refinement, resulting in a structure with the following particle count and nominal resolution: monomer RTC (456,629 particles; 2.68 Å) (Extended Data Fig. 4e). Local resolution calculations were generated using blocres and blocfilt from the Bsoft package ^20^ (Extended Data Fig. 4c). The angular distribution of particle orientations (Extended Data Fig. 4b) and directional resolution through the 3DFSC package ^41^ (Extended Data Fig. 4d) were calculated for the final class.

### S5_CTP

Grids were imaged using a CS corrected 300 kV Titan Krios (Thermo Fisher Scientific) equipped with a GIF BioQuantum and K3 camera (Gatan). Images were recorded with SerialEM ^43^ with a pixel size of 0.515 Å/px (micrograph dimension of 5760 × 4092 px) over a defocus range of −0.8 μm to −3.0 μm with a 20 eV energy filter slit. Movies were recorded in “counting mode” (native K3 camera binning 2) with ∼25 e-/px/s in dose-fractionation mode with subframes of 50 ms over a 2.5 s exposure (50 frames) to give a total dose of ∼57 e-/Å^2^. Dose-fractionated movies were gain-normalized, drift-corrected, summed, and dose-weighted using MotionCor2 ^37^. The CTF was estimated for each summed image using the Patch CTF module in cryoSPARC v3.1.0 ^38^. Particles were picked and extracted from the dose-weighted images with box size of 512 px using cryoSPARC Blob Picker and Particle Extraction. The entire dataset consisted of 13,905 motion-corrected images with 1,986,527 particles. Particles were sorted using three rounds of cryoSPARC 2D classification (N=50), resulting in 460,232 curated particles. An initial model (denoted as monomer) was generated using cryoSPARC *Ab initio* Reconstruction (N=3) on a subset of the particles. Particles were further curated using these initial models as 3D templates for iterative cryoSPARC Heterogeneous Refinement (N=6), resulting in 143,110 particles in the monomer class (cyan map, Extended Data Fig. 5) and 93,597 particles in an extracted dimer class (pink map, Extended Data Fig. 5). Curated particles in the monomer and dimer classes were re-extracted to a box size of 768 and input to cryoSPARC Homogenous and Non-uniform refinements ^39^. Particles within each class were further processed through two rounds of RELION 3.1 Bayesian Polishing ^40^. Polished particles were refined using cryoSPARC Local and Global CTF Refinement in combination with cryoSPARC Non-uniform Refinement, resulting in structures with the following particle counts and nominal resolutions: monomer RTC (128,484 particles; 3.09 Å) & dimer RTC (83,555 particles; 3.69 Å). To facilitate model building of the RTC, particles from the dimer RTC class underwent masked particle subtraction in which both protomers were masked, subtracted from the full map, and combined with the original monomer RTC to yield a structure of the RTC at a nominal resolution of 2.67 Å from 171,107 particles (Extended Data Fig. 5e). This combined RTC class was further refined with masked cryoSPARC Local Refinement using masks around the RdRp and NiRAN active sites. Locally refined maps were combined into a RTC composite map using PHENIX ‘Combine Focused Maps’ to aid model building ^42^. Local resolution calculations were generated using blocres and blocfilt from the Bsoft package ^20^ (Extended Data Fig. 5c). The angular distribution of particle orientations (Extended Data Fig. 5b) and directional resolution through the 3DFSC package ^41^ (Extended Data Fig. 5d) are shown for the final class.

### Model building and refinement

An initial model of the RTC was derived from PDB 7RE1 ^17^. The models were manually fit into the cryo-EM density maps using Chimera ^44^ and rigid-body and real-space refined using Phenix real-space-refine ^42^. For real-space refinement, rigid body refinement was followed by all-atom and B-factor refinement with Ramachandran and secondary structure restraints. Models were inspected and modified in Coot 0.9.5 ^45^ and the refinement process was repeated iteratively.

### Native mass spectrometry (nMS) analysis

The RTC samples were initially reconstituted as described above in the following buffer: 20 mM HEPES pH 8.0, 80 mM K-Acetate, 5 mM MgCl_2_, 2 mM DTT. For the NTP pre-incorporation experiments, the replication-transcription complexes with RNA scaffolds containing either 3’-oxy p-RNA (RTC) or 3’-deoxy p-RNA (RTC*) were incubated with 300 μM NTP (ATP, RDV-TP or GTP) on ice for 2 min prior to buffer exchange. For the NDP incubation experiments, the RTC samples reconstituted with the respective RNA scaffolds were incubated with 2 mM nucleotide (ADP, RDV-DP, GDP or ATP) on ice for 2 min before buffer exchange. After nucleotide incubation, all samples were immediately buffer exchanged into an nMS-compatible solution (150 mM ammonium acetate, pH 7.5, 0.01% Tween-20) with a 40 kDa MWCO (ThermoFisher Scientific). For nMS analysis, a 2 – 3 μL aliquot of the buffer-exchanged sample was loaded into a gold-coated quartz capillary tip prepared in-house and then electrosprayed into an Exactive Plus with extended mass range (EMR) instrument (Thermo Fisher Scientific) with a static direct infusion nanospray source ^46^. The MS parameters used include: spray voltage, 1.22 kV; capillary temperature, 125 - 150 °C; in-source dissociation, 0 - 10 V; S-lens RF level, 200; resolving power, 8,750 or 17,500 at *m/z* of 200; AGC target, 1 × 10^6^; maximum injection time, 200 ms; number of microscans, 5; injection flatapole, 8 V; interflatapole, 7 V; bent flatapole, 4 V; high energy collision dissociation (HCD), 200 V; ultrahigh vacuum pressure, 5.8 – 6.1 × 10^−10^ mbar; total number of scans, at least 100. Mass calibration in positive EMR mode was performed using cesium iodide. For data processing, the acquired MS spectra were visualized using Thermo Xcalibur Qual Browser (v. 4.2.47). Deconvolution was performed either manually or using the software UniDec v. 4.2.0 ^47,48^. The following parameters were used for data processing with UniDec: background subtraction (if applied), subtract curve 10; smooth charge state distribution, enabled; peak shape function, Gaussian. Mass accuracies were calculated as the % difference between the measured and expected masses relative to the expected mass. The observed mass accuracies (calculated as the % difference between the measured and expected masses relative to the expected mass) ranged from 0.005 – 0.06%.

The expected masses for the component proteins are nsp7: 9,137 Da; nsp8 (N-terminal Met lost): 21,881 Da, and nsp12 (has two Zn^2+^ ions coordinated with 6 deprotonated cysteine residues): 106,785 Da ^14^. The RNA scaffolds were also analyzed separately, and their sequences were verified by mass measurements using nMS.

### In-vitro primer elongation assays

Assays were performed using reconstituted template-primer RNA scaffolds (Table S1) (Horizon Discover Ltd./Dharmacon) annealed in 10 mM HEPES pH 8.0, 50 mM KCl, 2 mM MgCl_2_. Reactions (20 μL) containing 100 nM RNA scaffold, 0.75 μM nsp12, 2 μM nsp7/8 and NTPs 300 μM (if present as natural nucleotides, NDPs or α-β analogues) and 1 μL α-^32^P-GTP (Perkin-Elmer) were incubated at 30°C for 10 minutes prior to addition of a 2x stop solution (Invitrogen-Gel Loading buffer II). Assay buffer was 100 mM K-acetate, 20 mM HEPES pH 8.0, 5 mM MgCl_2 &_ 2 mM BME in which the MgCl_2_ was substituted with CaCl_2_ when monitoring the effects of alternative metal ions. Products of the elongation reactions were separated on 10% acrylamide-8M urea denaturing gels and analyzed by phosphorimaging.

### Quantification and statistical analysis

The local resolution of the cryo-EM maps (Extended Data Figs. 1-5) was estimated using blocres ^20^ with the following parameters: box size 15 & sampling with respective map pixel size. Directional 3DFSCs (Extended Data Figs. 1-5) were calculated using 3DFSC ^41^. The quantification and statistical analyses for model refinement and validation were generated using MolProbity ^49^ and PHENIX ^50^.

## Data and code availability

All unique/stable reagents generated in this study are available without restriction from the corresponding authors, Seth A. Darst and E.A. Campbell. The cryo-EM density maps and atomic coordinates have been deposited in the EMDataBank and Protein Data Bank as follows: S1_RDV-TP (EMD-26639, PDB 7UO4), S2_ATP (EMD-26641, PDB 7UO7), S3_UTP (EMD-26642, PDB 7UO9), S4_GTP (EMD-26645, 7UOB), S5_CTP (EMD-26646, PDB 7UOE).

## Acknowledgments

We thank Andreas Mueller and Ruth Saecker for helpful discussions. Some of the work reported here was conducted at the Simons Electron Microscopy Center (SEMC) and the National Resource for Automated Molecular Microscopy (NRAMM) and National Center for CryoEM Access and Training (NCCAT) located at the NYSBC, supported by grants from the NIH National Institute of General Medical Sciences (P41 GM103310), NYSTAR, the Simons Foundation (SF349247), the NIH Common Fund Transformative High Resolution Cryo-Electron Microscopy program (U24 GM129539) and NY State Assembly Majority. This work was supported by NIH P41 GM109824 and P41 GM103314 to B.T.C, and NIH R01 AI161278 and Gilead Sciences (to E.A.C. and S.A.D).

## Author contributions

B.F.M., J.C., J.K.P., T.K.A., J.Y.F., E.A.C. and S.A.D. conceived and designed this study. B.F.M. and J.C. performed cloning, protein purification and biochemistry. P.D.B.O. conducted mass spectrometry experiments. B.F.M. prepared cryo-EM specimens. Cryo-EM data were collected by B.F.M., E.C., J.M., E.T.E., M.E., J.S. and H.N. B.F.M. processed all cryo-EM data. B.F.M., J.K.P., E.A.C. and S.A.D. built and analyzed atomic models. E.A.C., R.L., B.T.C. and S.A.D supervised and acquired financial support. B.F.M. wrote the first draft of the manuscript; all authors contributed to the final version.

## Competing interests

E.A.C. and S.A.D. received funding from Gilead Sciences, Inc. in support of this study. J.K.P., T.K.A., J.Y.F., and J.P.B. are Gilead employees.

---

**Extended Data Fig 1.**
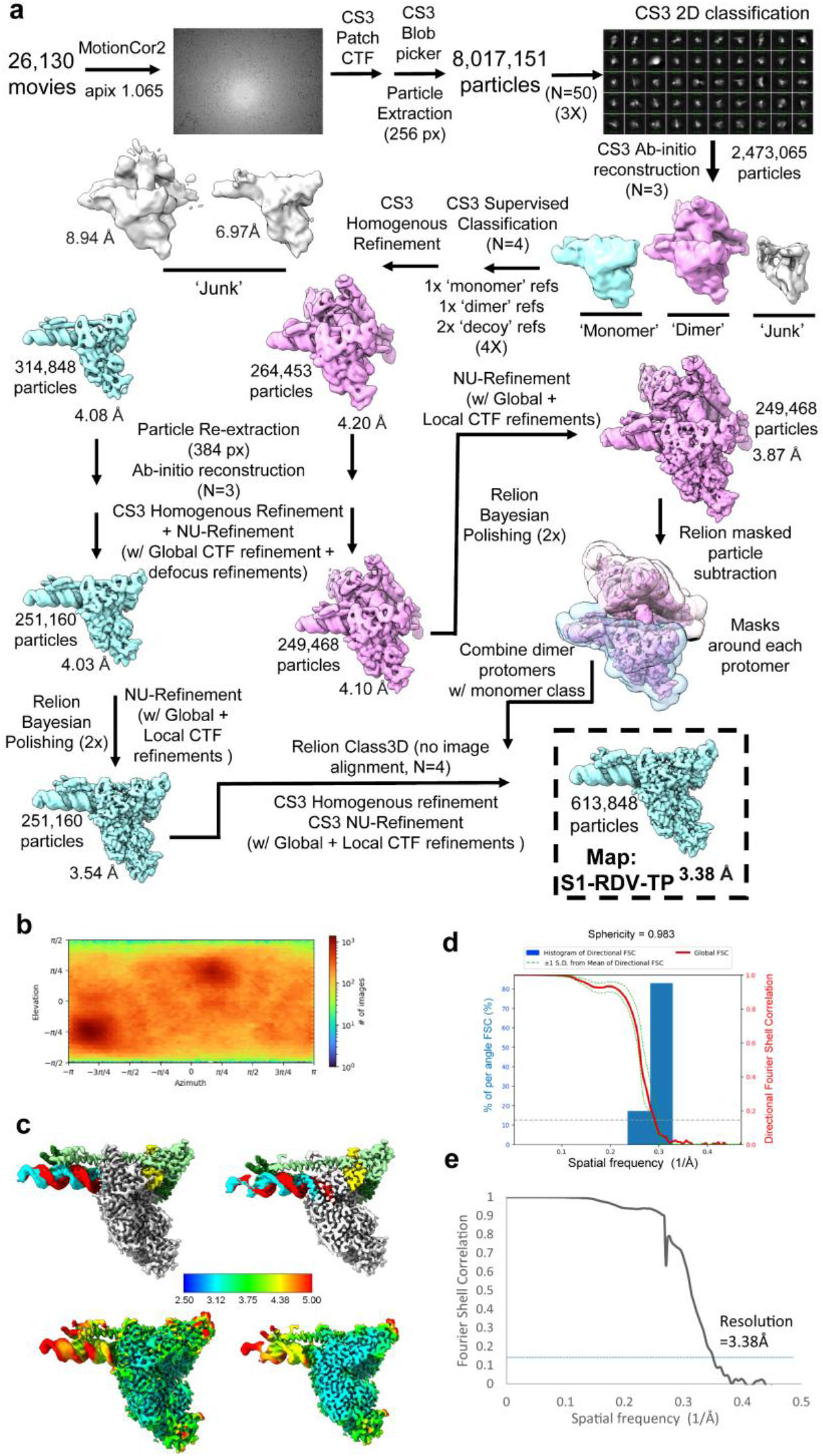
**(a)** Cryo-EM processing pipeline for the S1_RDV-TP dataset. **(b)** Angular distribution plot for the S1_RDV-TP dataset, calculated in cryoSPARC. Scale depicts number of particles assigned to a specific angular bin. **(c)** Nominal 3.38Å resolution cryo-EM reconstruction filtered by local resolution and colored according to fitted model chain. Right panel is clipped to reveal RTC active site. **(d)** Directional 3D FSC for S1_RDV-TP, determined with 3DFSC. **(e)** Gold-standard FSC plot for the S1_RDV-TP dataset, calculated by comparing two half maps from cryoSPARC. The blue dotted line represents the 0.143 FSC cutoff.

**Extended Data Fig 2.**
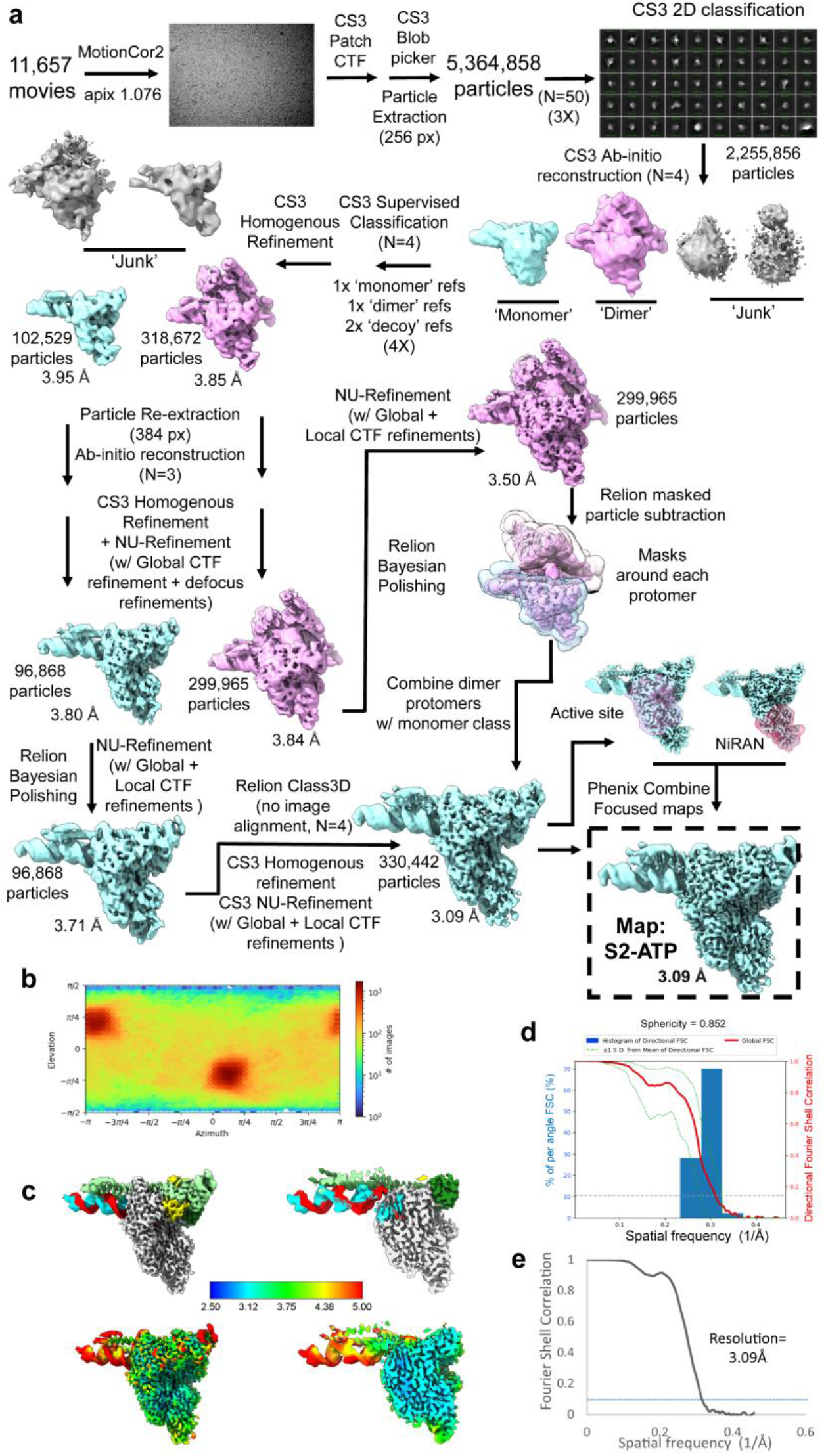
**(a)** Cryo-EM processing pipeline for the S2_ATP dataset. **(b)** Angular distribution plot for the S2_ATP dataset, calculated in cryoSPARC. Scale depicts number of particles assigned to a specific angular bin. **(c)** Nominal 3.09Å resolution cryo-EM reconstruction filtered by local resolution and colored according to fitted model chain. Right panel is clipped to reveal RTC active site. **(d)** Directional 3D FSC for S2_ATP, determined with 3DFSC. **(e)** Gold-standard FSC plot for the S2_ATP dataset, calculated by comparing two half maps from cryoSPARC. The blue dotted line represents the 0.143 FSC cutoff.

**Extended Data Fig 3.**
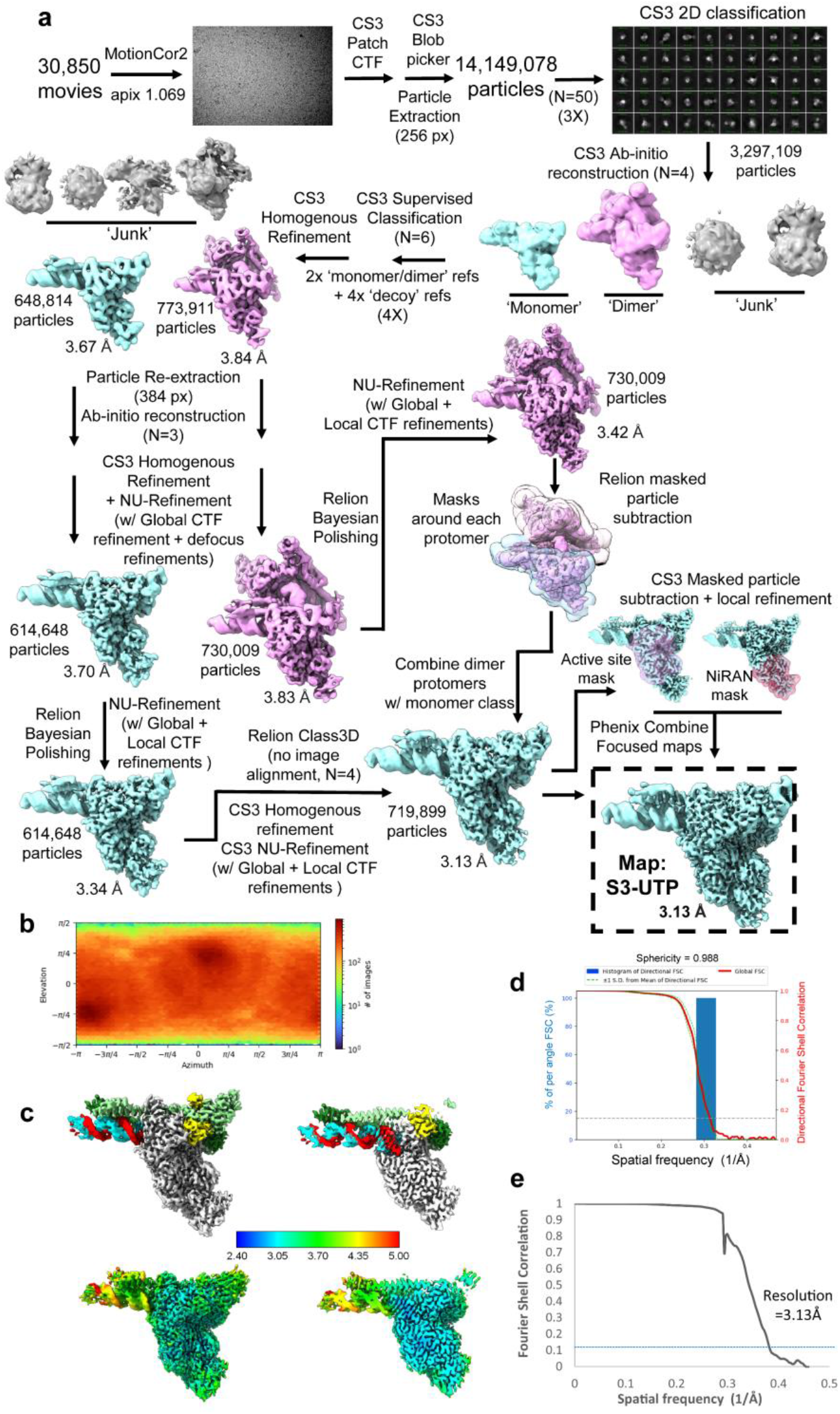
**(a)** Cryo-EM processing pipeline for the S3_UTP dataset. **(b)** Angular distribution plot for the S3_UTP dataset, calculated in cryoSPARC. Scale depicts number of particles assigned to a specific angular bin. **(c)** Nominal 3.13Å resolution cryo-EM reconstruction filtered by local resolution and colored according to fitted model chain. Right panel is clipped to reveal RTC active site. **(d)** Directional 3D FSC for S3_UTP, determined with 3DFSC. **(e)** Gold-standard FSC plot for the S3_UTP dataset, calculated by comparing two half maps from cryoSPARC. The blue dotted line represents the 0.143 FSC cutoff.

**Extended Data Fig 4.**
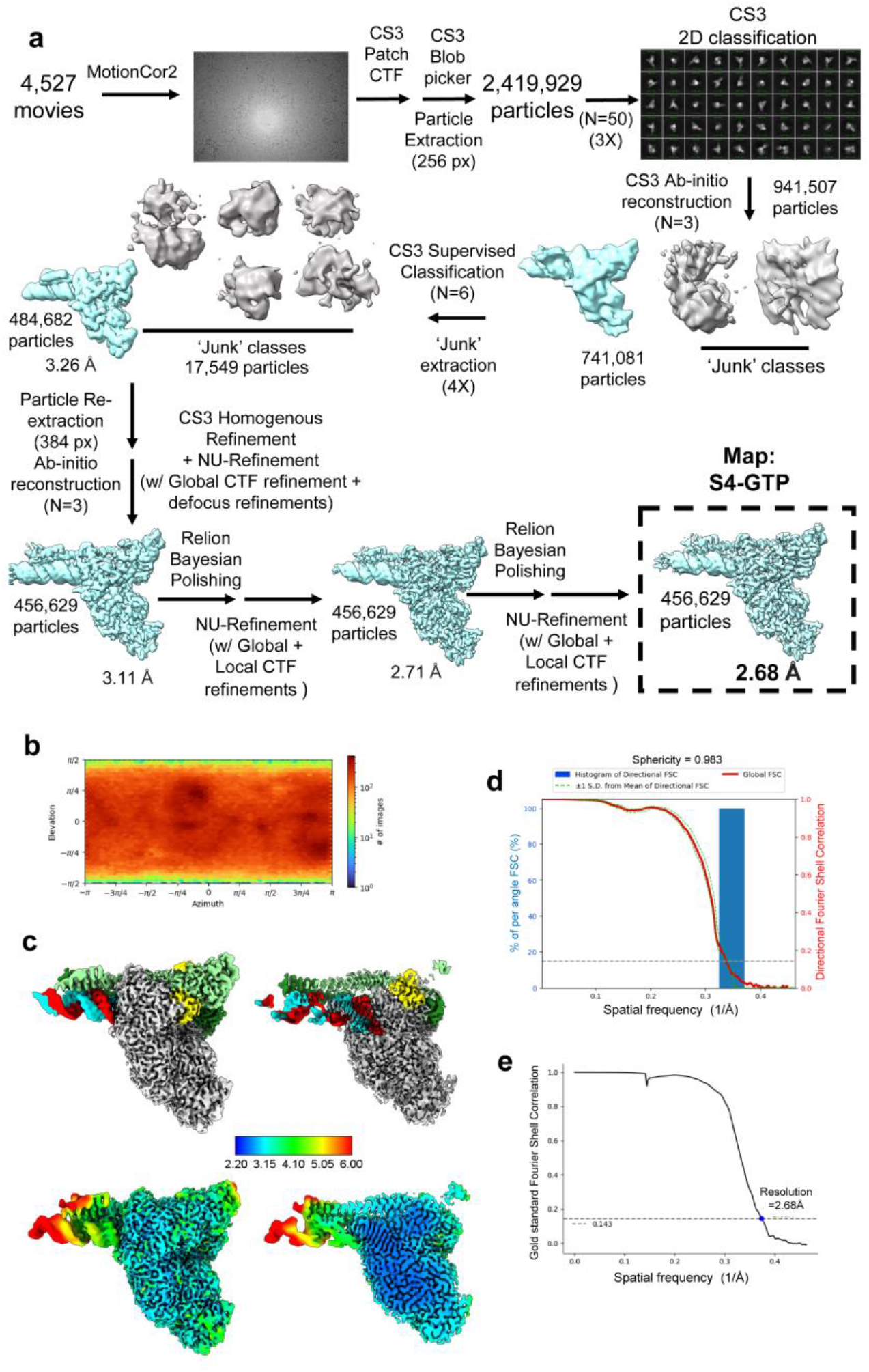
**(a)** Cryo-EM processing pipeline for the S4_GTP dataset. **(b)** Angular distribution plot for the S4_GTP dataset, calculated in cryoSPARC. Scale depicts number of particles assigned to a specific angular bin. **(c)** Nominal 2.68Å resolution cryo-EM reconstruction filtered by local resolution and colored according to fitted model chain. Right panel is clipped to reveal RTC active site. **(d)** Directional 3D FSC for S4_GTP, determined with 3DFSC. **(e)** Gold-standard FSC plot for the S4_GTP dataset, calculated by comparing two half maps from cryoSPARC. The black dotted line represents the 0.143 FSC cutoff.

**Extended Data Fig 5.**
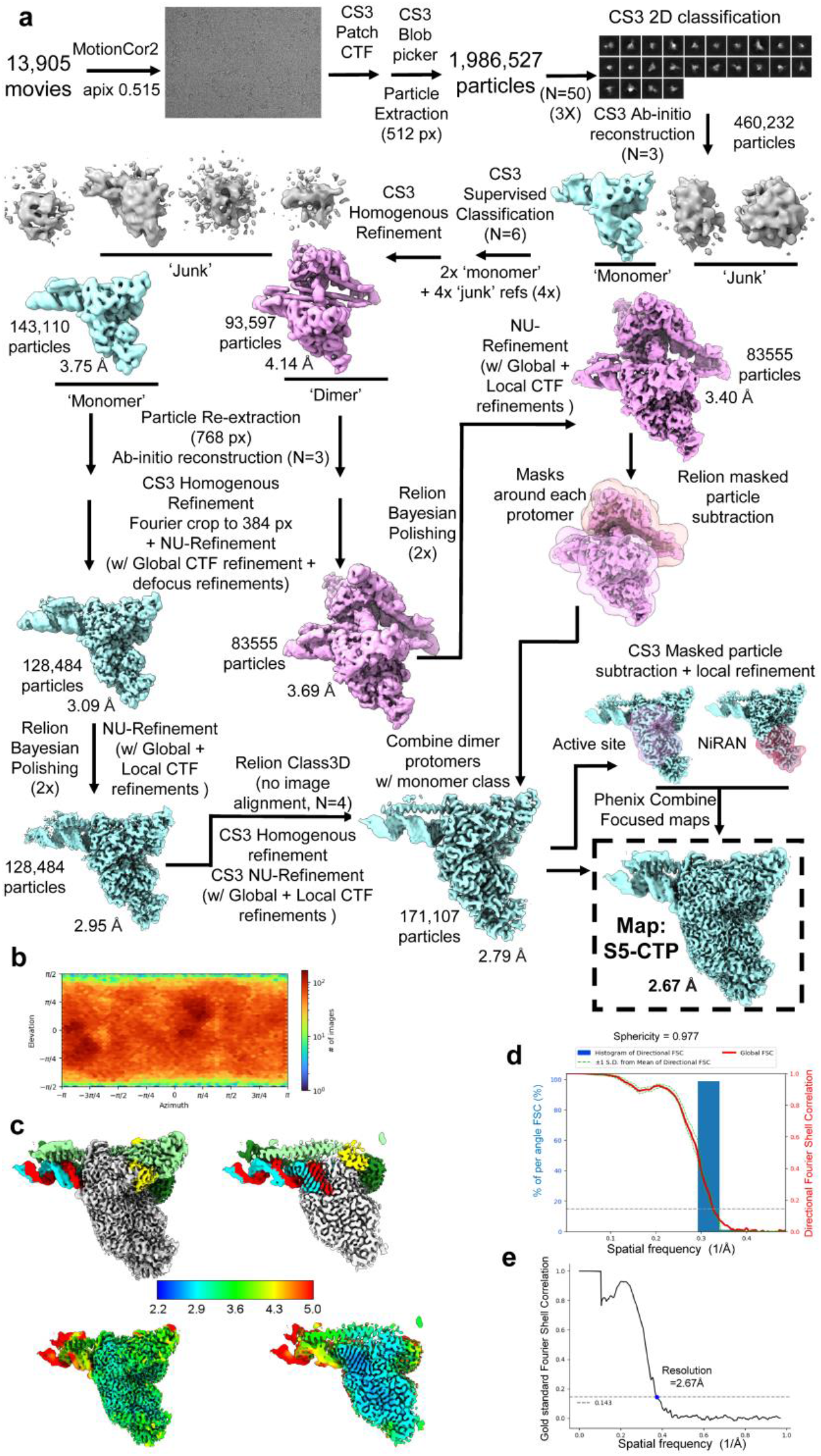
**(a)** Cryo-EM processing pipeline for the S5_CTP dataset. **(b)** Angular distribution plot for the S5_CTP dataset, calculated in cryoSPARC. Scale depicts number of particles assigned to a specific angular bin. **(c)** Nominal 2.67Å resolution cryo-EM reconstruction filtered by local resolution and colored according to fitted model chain. Right panel is clipped to reveal RTC active site. **(d)** Directional 3D FSC for S5_CTP, determined with 3DFSC. **(e)** Gold-standard FSC plot for the S5_CTP dataset, calculated by comparing two half maps from cryoSPARC. The black dotted line represents the 0.143 FSC cutoff.

**Extended Data Fig 6.**
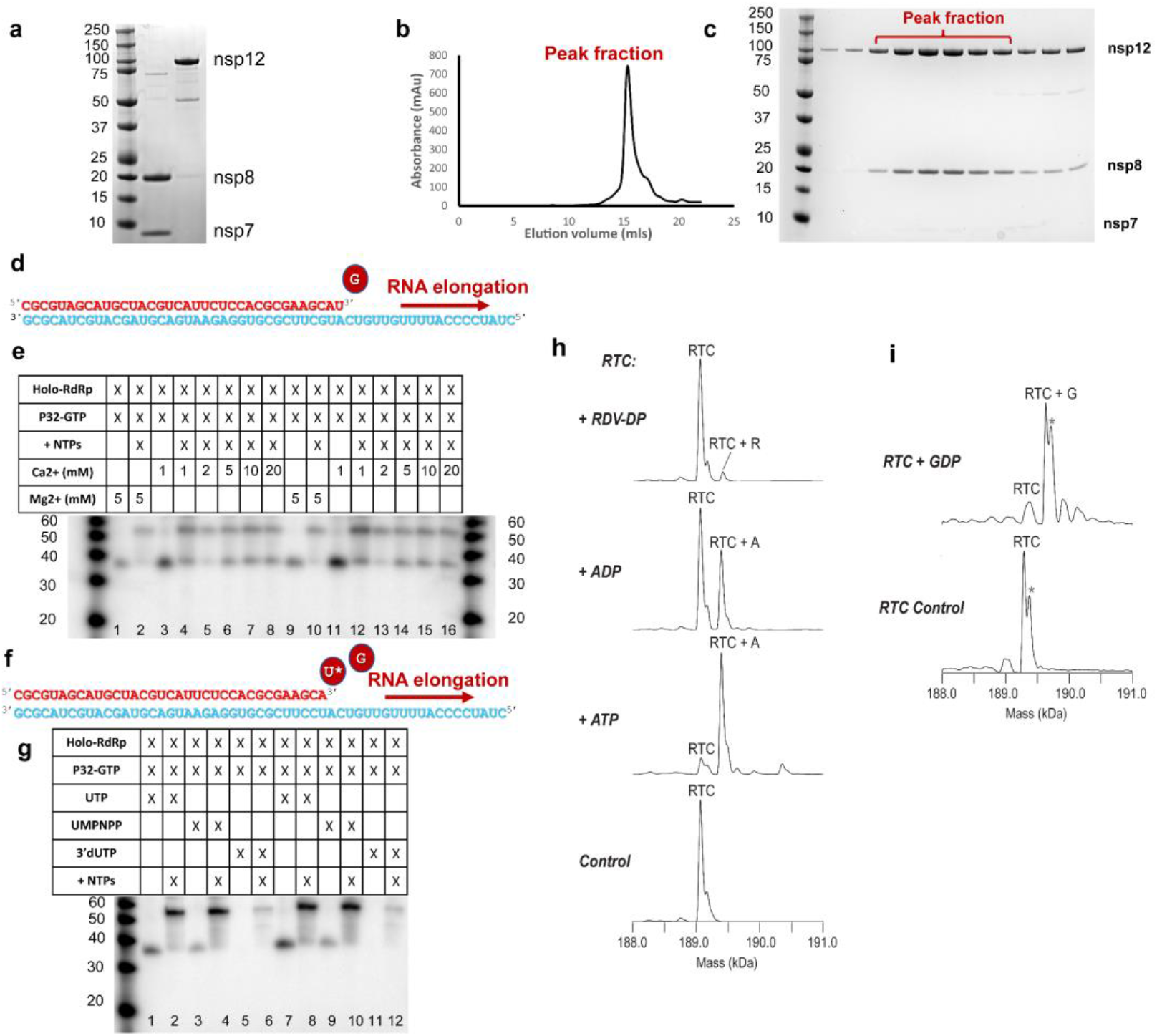
**(a)** SDS-PAGE of purified SARS-CoV-2 nsp7/8 & nsp12. **(b)** Size exclusion chromatography for the purified RTC complex, composed of nsp7/8_2_/12 bound to the reconstituted product/template-RNA scaffold. **(c)** SDS-PAGE of assembled RTC complex following size exclusion. **(d)** S4 RNA scaffold utilized for the S4_GTP structure and incorporation assays. **(e)** Gel-based primer elongation assay in presence of physiological metal, Mg^2+^, and non-physiological metal, Ca^2+^, to investigate use of Ca^2+^ for stabilization of the pre-incorporation complex (gel depicts n=2 with n=3 experiments performed). **(f)** S3 RNA scaffold used for the S3_UTP structure and incorporation assays. **(g)** Gel-based primer elongation assay in presence of UTP, UMPNPP and 3’deoxy UTP (gel depicts n=2 with n=3 experiments performed). **(h)** Native mass spectrometry analysis of RNA extension in presence of ADP, RDV-DP & ATP using the S1/S2 RNA scaffold. **(i)** Native mass spectrometry analysis of RNA extension in presence of GDP using the S4 RNA scaffold.

**Extended Data Fig. 7.**
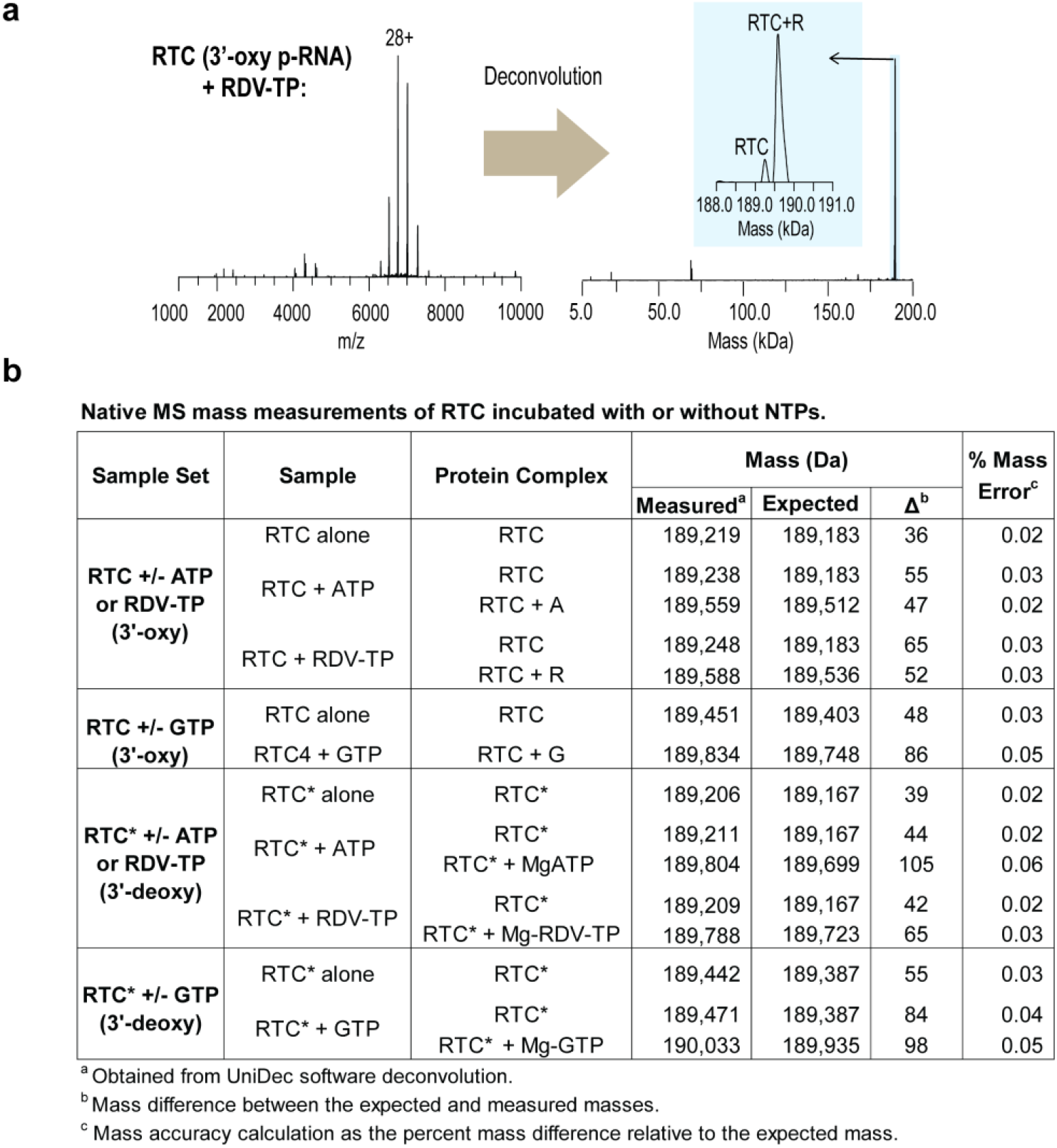
(**a**) Data processing and deconvolution of native MS spectra using the UniDec software. Analysis of a representative MS spectrum (sample: RTC + RDV-TP) is shown. (**b**) The table of native MS mass measurements obtained from UniDec deconvolution of the RTC and RTC* samples used in the NTP incorporation experiments

**Extended Data Fig 8.**
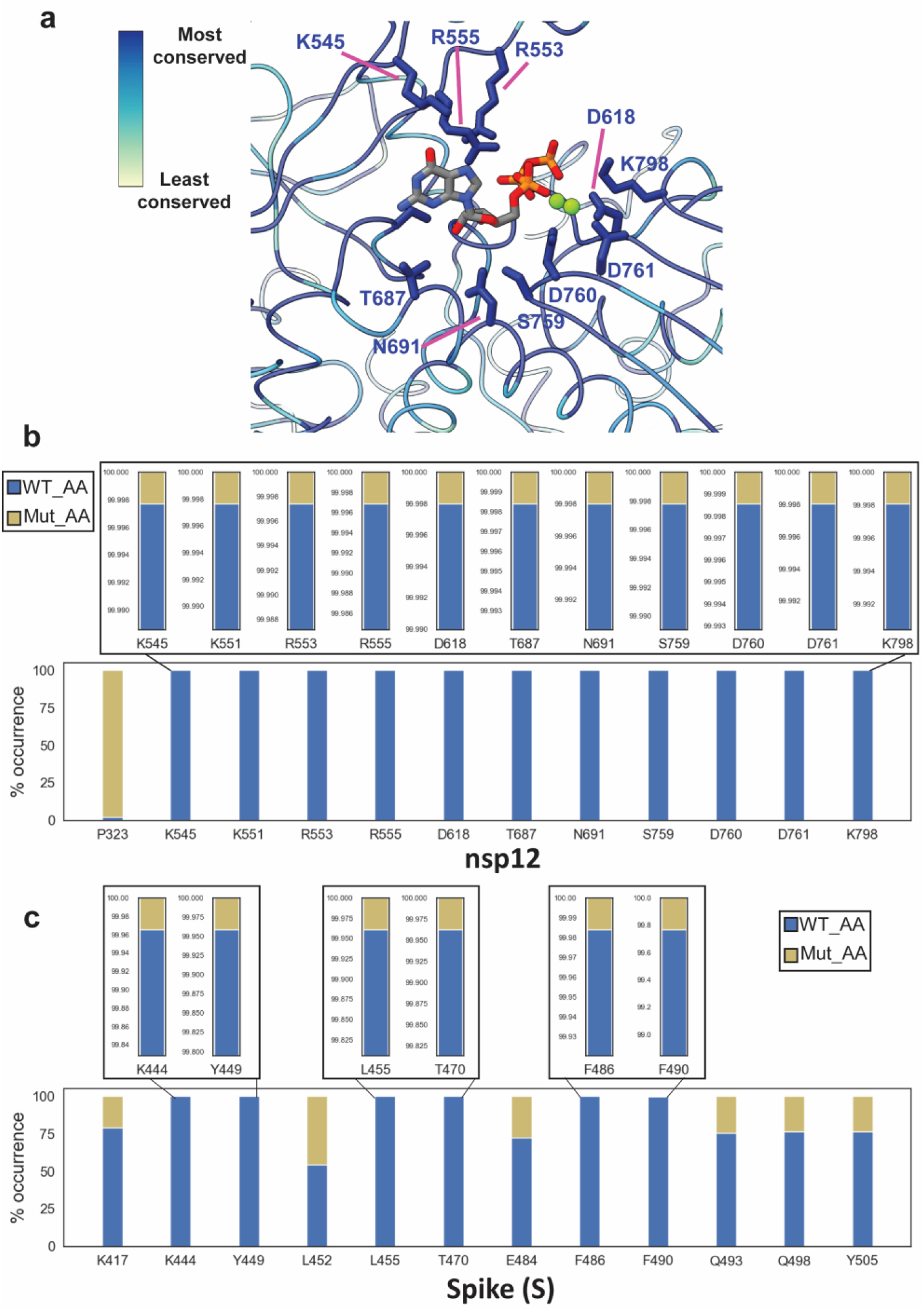
**(a)** Zoom-in on the active site of S4_GTP, highlighting residues (sticks) which interact with the bound GTP/2xMg in which ribbon/sticks are colored according to the amino acid conservation across a representative list of viruses found in the α & β coronavirus clades (Supplementary information). (b) Bar plot showing the frequency of occurrence of the wild-type amino acid (reference strain Wuhan/Hu-1/2019) as well as their summed mutations according to the GISAID database, as of April 2022, for the nsp12 active site residues and the nsp12 residue P323. (c) Bar plot showing the frequency of occurrence of the wild-type amino acid (reference strain Wuhan/Hu-1/2019) as well as their summed mutations according to the GISAID database, as of April 2022, for Spike (S) residues found in the ACE2 binding region.

**Extended Data Fig 9.**
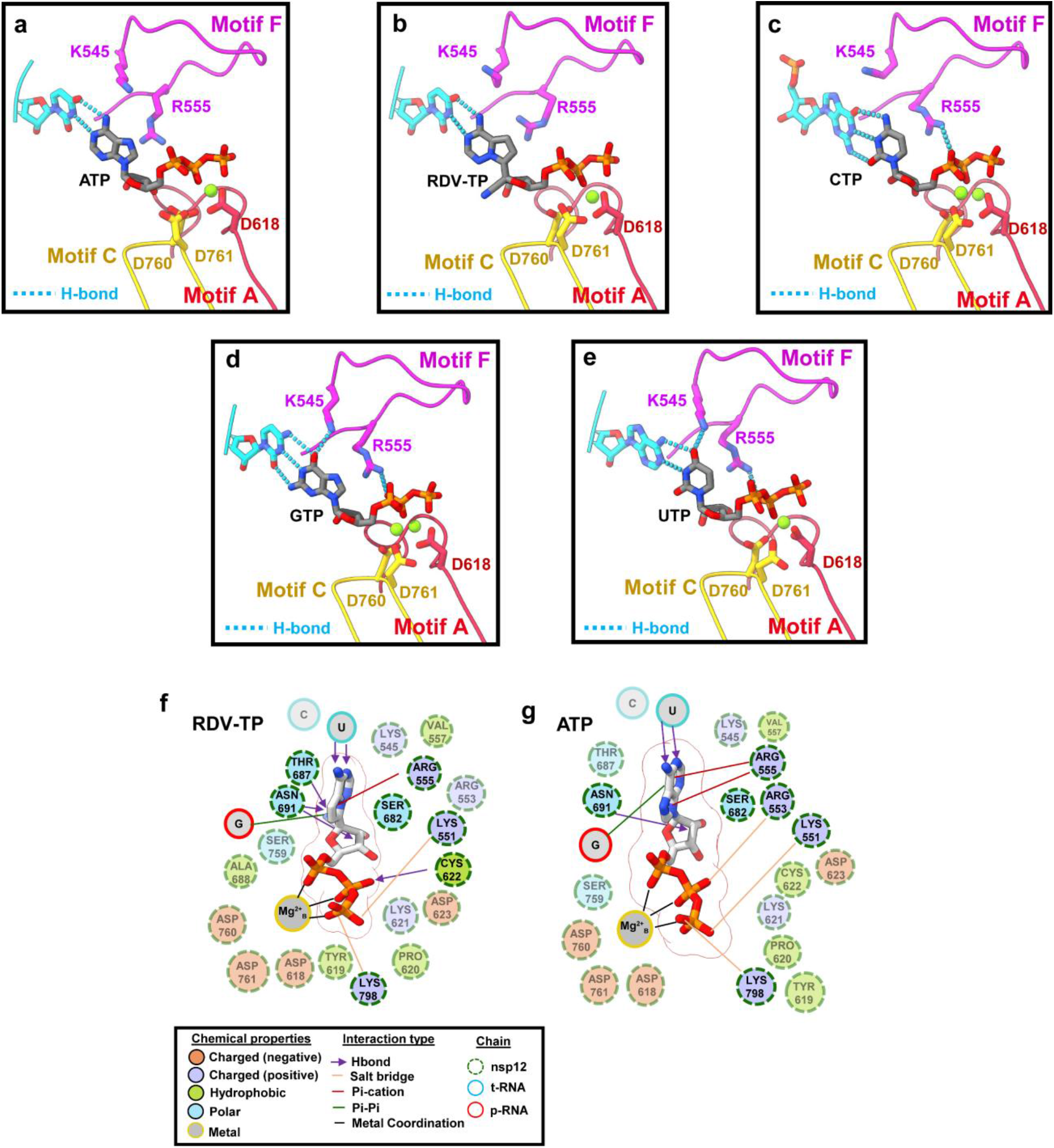
**(a - e)** Analysis of the active sites of structures (S1-S5) illustrate that K545 and R555 interact differentially with each of the incoming nucleotides. H-bonds are depicted as dotted-cyan lines. **(f, g)** 2D schematics illustrating the set of interactions between RDV-TP **(f)** and ATP **(g)** and the active site,

**Extended Data Fig 10.**
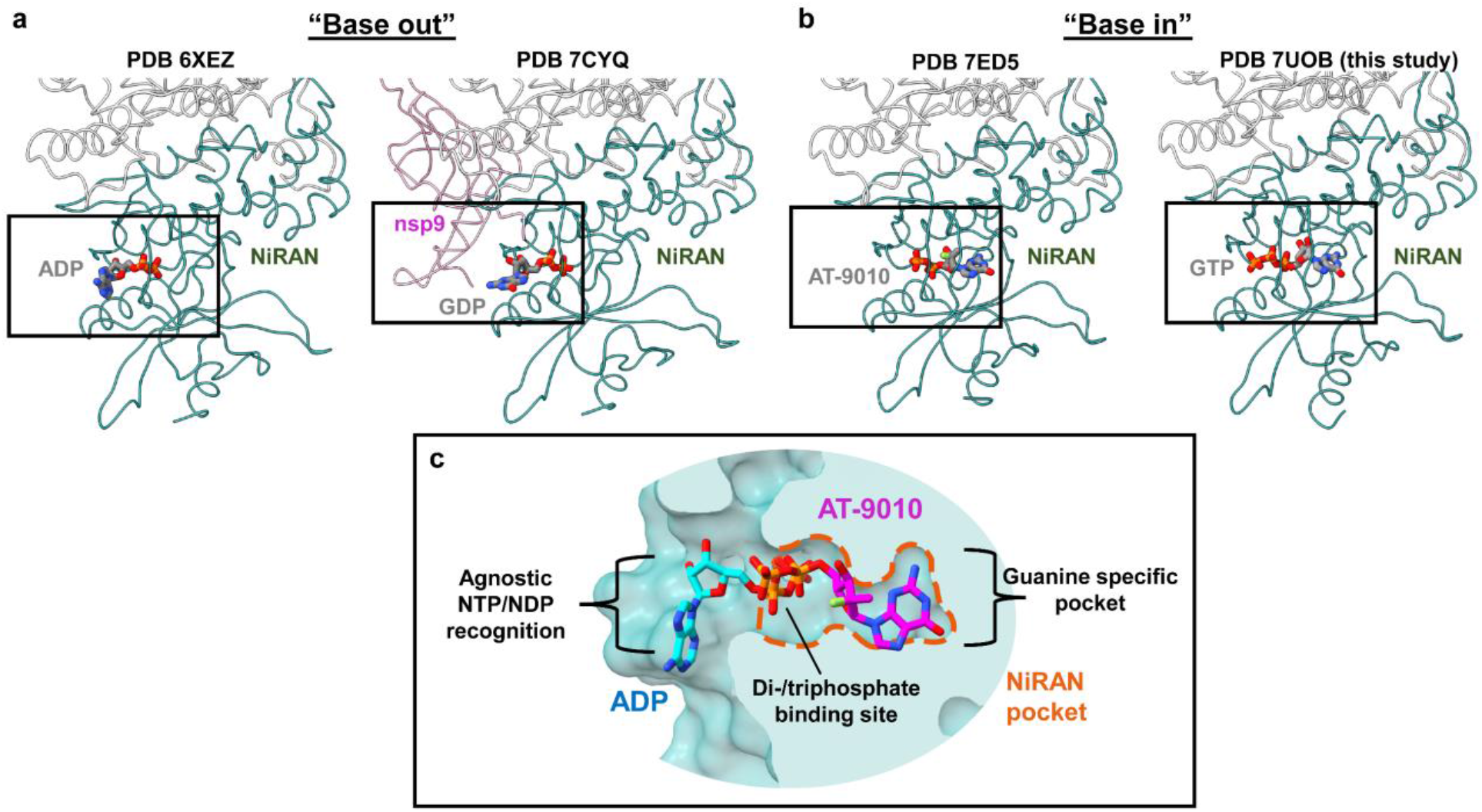
**(a)** Models (PDBs 6XEZ & 7CYQ) of the nsp12 NiRAN with a bound base in the ‘Base-out’ pose. (b) Models (PDBs 7ED5 & 7UOB) of the nsp12 NiRAN with a bound base in the ‘Base-in’ pose. (c) Surface representation detailing the NiRAN active site pocket bound to a nucleotide in the ‘base-out’ pose (PDB 6XEZ) aligned with a structure with a bound nucleotide in the ‘base-in’ pose (PDB 7ED5).

## Supplementary information

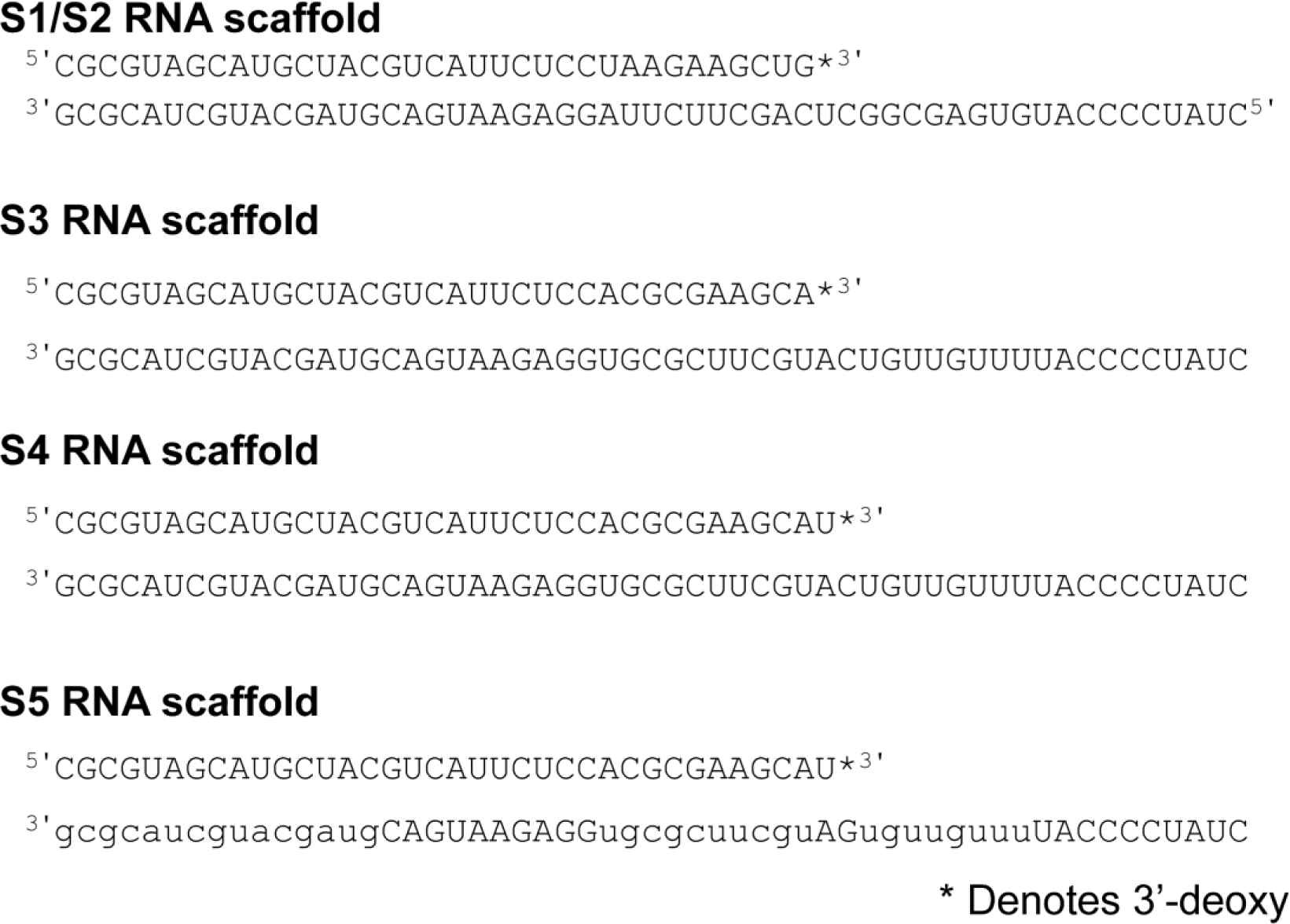

